# Sequence characterization and molecular modeling of clinically relevant variants of the SARS-CoV-2 main protease

**DOI:** 10.1101/2020.05.15.097493

**Authors:** Thomas J. Cross, Gemma R. Takahashi, Elizabeth M. Diessner, Marquise G. Crosby, Vesta Farahmand, Shannon Zhuang, Carter T. Butts, Rachel W. Martin

## Abstract

The SARS-CoV-2 main protease (M^*pro*^) is essential to viral replication and cleaves highly specific substrate sequences, making it an obvious target for inhibitor design. However, as for any virus, SARS-CoV-2 is subject to constant selection pressure, with new M^*pro*^ mutations arising over time. Identification and structural characterization of M^*pro*^ variants is thus critical for robust inhibitor design. Here we report sequence analysis, structure predictions, and molecular modeling for seventy-nine M^*pro*^ variants, constituting all clinically observed mutations in this protein as of April 29, 2020. Residue substitution is widely distributed, with some tendency toward larger and more hydrophobic residues. Modeling and protein structure network analysis suggest differences in cohesion and active site flexibility, revealing patterns in viral evolution that have relevance for drug discovery.

## Introduction

Severe acute respiratory syndrome coronavirus 2 (SARS-CoV-2) emerged in late 2019 (*1*) and rapidly spread worldwide, causing an ongoing pandemic. Although the sequence of its RNA genome is highly similar to that of SARS-CoV-1, SARS-CoV-2 is believed to have arisen independently from a bat coronavirus (*2*), to which it shares 96% similarity (*3*). The emerging SARS-CoV-2 subsequently gained a modified spike protein due to recombination in an intermediate host, the pangolin (*4, 5*), followed by purifying selection for binding to the human ACE2 protein (*6*). No therapeutic agents able to reduce SARS-CoV-2 mortality in clinical settings are yet known, although extensive efforts are underway to discover new drugs or repurpose existing ones to inhibit key viral proteins. Here we focus on the main protease (M^*pro*^), which plays a critical role in viral replication. Like other betacoronaviruses, SARS-CoV-2 is a positive-sense RNA virus that expresses all of its proteins as a single polypeptide chain, which is cleaved by M^*pro*^ to yield the mature proteins (*7*).

Inhibiting this key enzyme would prevent viral replication, reducing viral load and thus symptom intensity. A similar approach was instrumental in making HIV a manageable disease (*8–10*). However, the proteins in question differ markedly, rendering HIV protease inhibitors ineffective against SARS-CoV-2; indeed, a standard HIV protease inhibitor combination did not prove effective against COVID-19 in a recent clinical trial (*11*). Specifically, HIV protease is an aspartic protease (and functional only as a dimer, as the active site comprises one residue from each monomer), whereas M^*pro*^ is a 3CL cysteine protease that is likewise most active in the dimeric state, although each monomer has its own catalytic dyad (*12*). The 3CL cysteine proteases are characterized by a chymotrypsin-like fold and a cysteine-histidine catalytic dyad in the active site, implying both different structures and distinct chemical mechanisms. While the general strategy of seeking protease inhibitors is hence viable for both SARS-CoV-2 and HIV, drug development for the former depends on characterizing this novel enzyme.

Molecular modeling is an important tool for guiding inhibitor discovery, making it possible to evaluate large numbers of candidate drugs *in silico* to select experimental targets; however, standard approaches screen against only one version of the protein, typically the reference or wild-type (WT) sequence. In a host population, mutations accumulate with each viral passage, generating a *mutational landscape* rather than a single protein. The design of robust inhibitors that can protect against the multiple strains encountered in clinical settings requires characterization of this sequence space and the populations of conformations it engenders. Furthermore, effective and rapid response to future emerging coronavirus diseases requires both *in silico* screening and experimental testing of antiviral agents and a validated library of relatively general inhibitors that can be used as a basis for the development of specialized therapeutics. Central to the success of that effort will be developing an understanding of structural and functional variation in SARS-CoV-2 proteins, particularly as mutations accumulate and new strains emerge. Here we characterize all 79 known variants of M^*pro*^ as of 29 April, 2020, and analyze trends in amino acid substitutions and the resulting structural changes using network analysis and molecular modeling. To our knowledge this is the first detailed analysis of clinically relevant mutations in M^*pro*^. Our analysis shows a trend toward substitution for larger and more hydrophobic residues versus the WT protein. Analysis of active site networks (ASN) from M^*pro*^ variants suggests differences in active site flexibility and cohesion that may serve to guide the design of robust, mutation-resistant inhibitors.

## Results and Discussion

### Mutations in M^*pro*^ are geographically distributed and occur throughout the protein

From the GISAID (https://www.gisaid.org/) (*13*) EpiCoV database (through 29 Apr, 2020), 78 unique non-synonymous mutations to M^*pro*^ were found in addition to the WT sequence, including 73 single point variants and 5 double variants. For genome sequences containing these M^*pro*^ variants, full genome alignments were performed using MUSCLE (*14*), and neighbor-joining trees were generated using MEGA X (*15*). Overall, the variation in SARS-CoV-2 sequences observed so far is relatively low, with mutation hotspots not evenly distributed throughout the genome, but localized to specific sequence regions (*16*). Because M^*pro*^ is critical for viral replication, mutations that have a large deleterious effect on virus replication are unlikely to be observed in clinical isolates; all M^*pro*^ variants investigated here are therefore assumed to be enzymatically competent. In general, codon usage and amino acid frequency in viruses of eukaryotes are essentially identical to those of their eukaryotic hosts, reflecting the viruses’ use of the host translation machinery (*17*). M^*pro*^ sequences found in sequences isolated from human hosts will therefore likely reflect bias toward human codon usage, somewhat limiting the scope of the observed mutation space.

The known mutations in M^*pro*^ are summarized in Figure 1. The tree was generated based on overall genome similarity; however, only sequences containing at least one mutation in M^*pro*^ were included in the analysis, along with the WT human sequence and two non-human reference sequences. The accession numbers and geographical sources are listed in Supplementary Table S2. The solid arcs around the outside of the diagram indicate M^*pro*^ mutations; color coding corresponds to the geographical source. Several mutations appear to have arisen more than once in the virus’s evolutionary history so far. Notably, K90R variants appear in multiple distantly related subtrees; five of these unique evolutionary events can be verified in Nextstrain’s SARS-CoV-2 phylogenetic tree (*18*). Further, L89F, P108S, and N274D arise at least twice in both trees.

**Figure 1:**
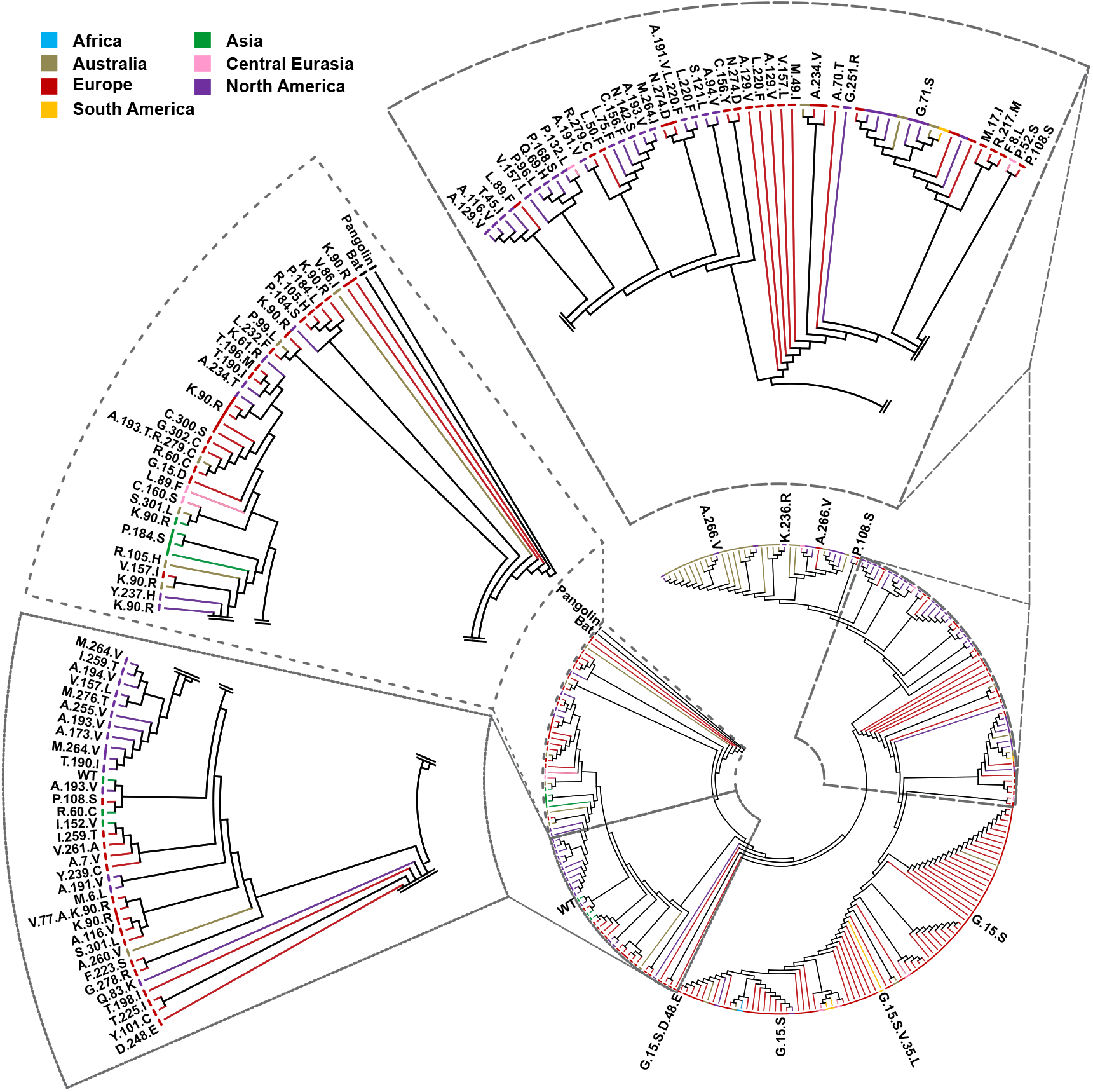
Optimal tree generated using 512 full mutant genomes and three reference genomes: human wild-type (WT) (*22*), bat (*3*), and pangolin (*19*). Only topology is shown; branch lengths are not to scale (average branch length = 1.432161×10^−4^ base substitutions per site). Each continuous arc corresponds to a variant label; these represent only adjacent branches with the same mutation in M^*pro*^, and do not necessarily indicate shared ancestry. Branches and arcs from human clinical samples are color coded by location, which includes the following subregions: **Africa**, light blue (Democratic Republic of the Congo); **Asia**, green (Beijing, Fujian, Malaysia, Shanghai, Vietnam, and Wuhan); **Australia**, gold; **Central Eurasia**, pink (Georgia, Jordan, Russia, and Turkey); **Europe**, red (Belgium, Denmark, England, Finland, France, Germany, Iceland, Luxembourg, Netherlands, Scotland, Spain, Sweden, Switzerland, and Wales); **North America**, purple (Costa Rica and United States of America); **South America**, yellow (Argentina and Brazil). Subtrees that contained identical subregions and mutations have been condensed into a single branch; all subtrees and their constituent accessions can be found in Supplementary Table S2.

These phylogenetic comparisons appear to support a multiple event hypothesis, but are subject to errors resulting from the sparsity of testing. The repeated occurrence of the same mutation in seemingly unrelated subtrees may be due to missing data that would show their evolutionary connectedness. The average branch length of Figure 1, which shows only topology, is 1.432161×10^−4^ base substitutions per site (including those from the bat (*3*) and pangolin (*19*)); 32.2% of the 1028 branches have, to ten significant figures, 0 base substitutions per site. For a genome of roughly 30,000 base pairs, this amounts to an average of only 4 substitutions per branch. All of these unique mutants therefore effectively belong to the same strain, making them difficult to place in an evolutionary context. For more diverged mutants, unfortunately-placed ambiguous nucleotides (*20*) could push them from one subtree to another. With the exception of five double variants, a majority of the sequences in Figure 1 arise from single point mutations. Whether and how M^*pro*^ mutations have affected viral fitness is not yet known, but at least three mutants have remained in the population long enough to accumulate another mutation: L220F to A191V/L220F, G15S to G15S/D48E and G15S/V35L, and K90R to V77A/K90R. It is worth noting that although a single variant A191V exists, the A191V/L220F double variant likely stemmed from an L220F ancestor due to its shared lineage with L220F single variants. A fifth double variant, A193T/R279C, was found but did not stem from any single mutation in our dataset; its origins remain unclear.

While a mutation’s prevalence and evolution in a population may be interpreted as a sign of stable viral function, the opposite does not necessarily indicate reduced virulence. Testing rates, social behavior, and time of first infection in each region are all factors that contribute to the spread of the disease and the availability of sequencing data. For instance, a large number of K90R mutants were collected in Iceland, where the number of tests per 1,000 people is nearly twice as many as the next leading country’s and more than seven times as many as the United States’ (Iceland: 141.75, USA: 18.21, as of 29 April, 2020) (*21*). Consequently, further investigation is needed to determine whether M^*pro*^ mutations affect viral fitness on a global scale. As such, without greater divergence and more sequences, it is difficult to tell if the presence of an M^*pro*^ mutation in unrelated subtrees is evidence of multiple evolutionary events, or an artifact of sparse testing.

Because only sequences harboring M^*pro*^ mutations were retained for analysis, certain geographical areas appear to be underrepresented. It is likely that the strains that had spread to underrepresented regions prior to our data collection simply did not have M^*pro*^ mutations. Different regions tend to be dominated by different mutants, a feature that might be explained by the timing at which these mutations arose or arrived. For instance, 83 of the 100 M^*pro*^ mutants from Iceland were K90R, and most stemmed from a single shared ancestor (Supplementary Table S2). Further, it is likely that heterogeneity in sequencing rates have resulted in a less-than-complete dataset. As of April 29th, the only North American, South American, and African M^*pro*^ mutants reported in the GISAID database that passed our filtering parameters were from Costa Rica and the USA, Argentina and Brazil, and the DRC respectively. This does not necessarily indicate a lack of M^*pro*^ mutations in other subregions, and may instead reflect differences in sequencing rates. In the structural analyses that follow, we focus on the differences in protein properties relative to WT, of the clinically observed M^*pro*^ variants.

### M^*pro*^ mutations to date suggest selection for larger, more massive, and more hydrophobic residues

To reveal the global pattern of substitutions, we visualize mutations in M^*pro*^ - independent of sequence position or location in the three-dimensional structure - by a network in which the nodes, or vertices, are amino acid types and the edges (represented by arrows pointing in the direction of substitution) are directional indicators of how often one amino acid was observed to substitute for another (Figure 2). The weights of the edges indicate the frequency of the mutation across known M^*pro*^ variants, while node color reflects residue hydrophobicity on the scale of (*23*) (larger numbers indicate greater hydrophobicity.) The most obvious trend observed in the pattern of mutation so far is the preferential substitution of larger, more hydrophobic amino acids in place of smaller, less hydrophobic ones. Overall, the pattern is consistent with increased incidence of amino acids that are more likely to be present in folded domains, rather than in linker regions (*24*).

**Figure 2:**
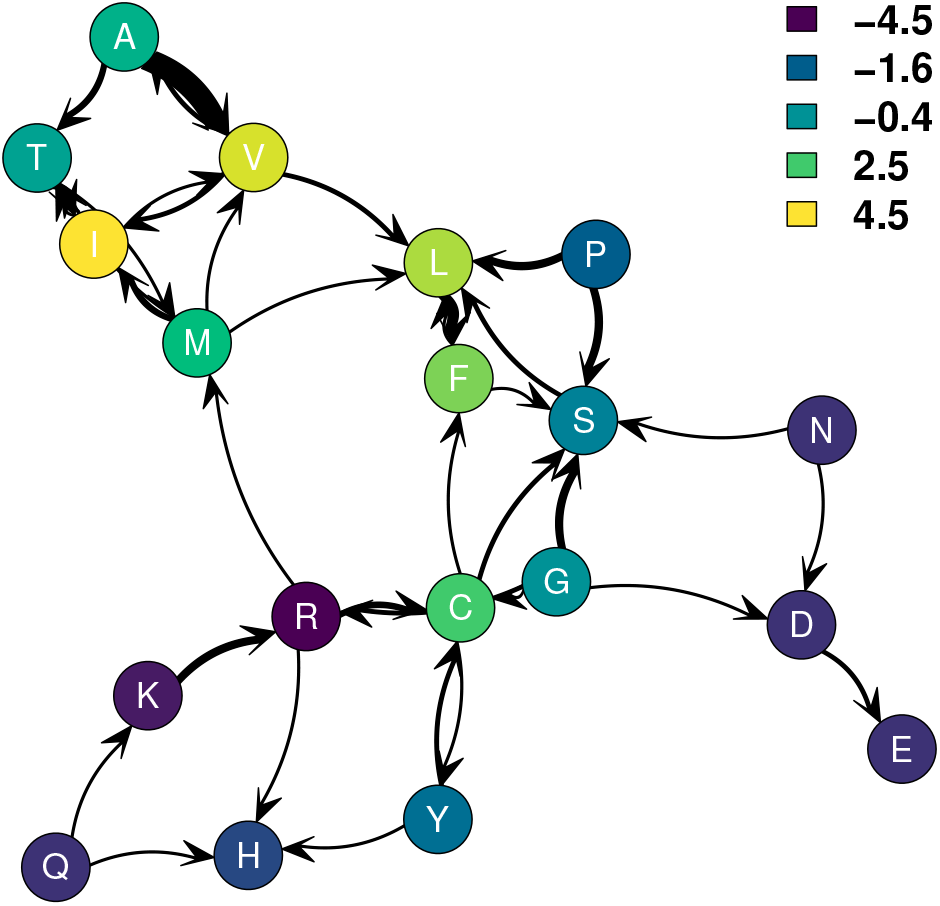
Amino acid substitutions observed to date in SARS-CoV-2 M^*pro*^. Arrows indicate direction of substitution: an arrow from *i* to *j* indicates at least one clinically observed substitution of residue type *i* to residue type *j*; heavier lines indicate larger numbers of observed substitutions. Color indicates hydrophobicity, using the scale of Kyte and Doolittle (*23*). In general, substitution has been towards larger and more hydrophobic residues.

In particular, it is notable that alanine has very few incoming ties and a large number of outgoing ties, mostly to valine, which has a larger and more hydrophobic side chain. Alanine is at the same time one of the most common amino acids and one of those with the most variable prevalence in the human genome (*25*). Similarly, observed ties to isoleucine are mostly incoming from smaller residues, and leucine, which is also large and hydrophobic, likewise has more incoming than outgoing ties overall, with the bulk of its outgoing ties going to phenylalanine. However, aromatic residues per se do not appear to be selected at a higher rate than can be explained by their hydrophobicity. Also notable is the selection away from the secondary structure-breakers proline and glycine, both of which have only outgoing ties, and the propensity for lysine to be replaced by arginine even though both side chains are positively charged. Arginine is both larger and capable of making more and stronger hydrogen bonds, as well as cation-*π* interactions not available to lysine, leading to its known overrepresentation in inter-domain and inter-monomer interfaces (*26–29*).

The mean differences in sidechain properties for observed M^*pro*^ mutations are summarized in Table 1. As observed in the network representation (Figure 2), mutated residues are, on average, larger and more hydrophobic than those they replace. Although substituted residues are on average larger and more massive, we do not see strong evidence favoring bulky over compact residues net of mass: residue bulk (measured as volume/mass) for substituted residues did not differ significantly from WT (mean difference=0.02Å ^3^/Da, *t* = 1.87, *p* = 0.0650). The variant sequences are not significantly different from WT in charge or aromatic content.

**Table 1:** Mean differences in side chain properties for substituted residues, versus WT (*N* = 83; substitutions from double mutants considered separately). On average, substituted residues are significantly more hydrophobic, massive, and larger than those they replace (all *p*-values for two-tailed *t*-tests versus no difference).

| | Mean Difference | Std. Err | $t$ value | $p$-value | |
| --- | --- | --- | --- | --- | --- |
| Polar (1=True) | 0.08 | 0.07 | 1.22 | 0.2251 |  |
| Hydrophobicity | 1.03 | 0.30 | 3.47 | 0.0008 | *** |
| Charge | -0.05 | 0.04 | -1.27 | 0.2078 |  |
| Aromatic (1=True) | 0.07 | 0.04 | 1.62 | 0.1093 |  |
| Mass (Da) | 9.97 | 3.49 | 2.85 | 0.0055 | ** |
| Volume (Å<sup>3</sup>) | 11.58 | 3.65 | 3.17 | 0.0021 | ** |
| Bulk (Å<sup>3</sup>/Da) | 0.02 | 0.01 | 1.87 | 0.0650 |  |
| Sig. codes: * $p < 0.05$, ** $p < 0.01$, *** $p < 0.001$ | | | | | |

### Molecular modeling suggests regionally specific differences in M^*pro*^ variant structure

For WT and each M^*pro*^ variant, a molecular model was constructed using MODELLER 9.23 (*30*), based on the A chain monomer of the PDB structure 6Y2E (*31*), followed by annealing, correction of protonation states, and all-atom molecular dynamics simulation in explicit solvent (see Methods). Examples of representative models are shown in Supplementary Figure S1, with the positions of all mutated residues shown mapped onto the WT structure in Supplementary Figure S2. We do not observe gross differences in structure or dynamics across variants, as expected given that all variants were found in clinical isolates and are therefore necessarily functional; mutations leading to radically altered or misfolded structures would likely be strongly selected against. However, analysis of MD trajectories does suggest more subtle differences across variants, providing insight into function-preserving changes.

To assess the overall degree to which local structure is conserved across M^*pro*^ variants, we compute the cross-variant variance in average *ϕ, Ψ* backbone torsion angles by residue. In order to control for overall flexibility, we normalize this by the estimated variance in torsion angles within each trajectory. For arbitrary angle *α*_*i*_ at residue *i*, this leads to the *local variation index*

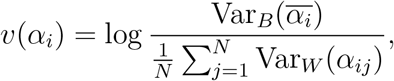

where *α*_*ij*_ is the vector of angles of type *α*_*i*_ over the trajectory of variant *j* with corresponding angular mean 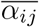, 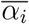 is the vector of such means across variants, Var_*B*_ is the “between variant” angular variance in mean angles, and Var_*W*_ is the “within variant” angular variance in *α*_*ij*_. Intuitively, high values of *v*(*α*_*i*_) indicate relatively large between-variant variation in *α*_*i*_ relative to angular variation seen within the trajectories themselves. For *v*(*ϕ*_*i*_) and *v*(*Ψ*_*i*_), such values correspond to systematic changes in local conformation associated with M^*pro*^ mutations. By turns, low values of *v*(*ϕ*_*i*_) and *v*(*Ψ*_*i*_) indicate residues whose local structure does not vary meaningfully across variants. It should be noted that such regions can be either flexible or rigid.

Figure 3 shows the mean local variation indices for *ϕ, Ψ* by residue for the 79 M^*pro*^ variants, indicated by color on the structure of M^*pro*^ WT. (Separate values for *ϕ* and *Ψ* are shown in Supplementary Figure S3.) It is immediately noteworthy that - with the minor exception of two small loop regions around N277 and F223 (respectively) - domain 3 shows little systematic variation across variants. The *β*-sheet-rich structure around the active site is also relatively well conserved. By contrast, we see relatively high levels of between-variant difference in the inter-domain region involving the termini (residues G2-A7 and S301-F305) and the double loop “active site gateway” region involving (respectively) L50-Y54 and D187-A191. The former is potentially significant in influencing large-scale flexibility (possibly relevant to dimerization), whereas the latter is of obvious relevance to substrate processing and specificity. This motivates a more detailed examination of variation in the active site, to which we return below.

**Figure 3:**
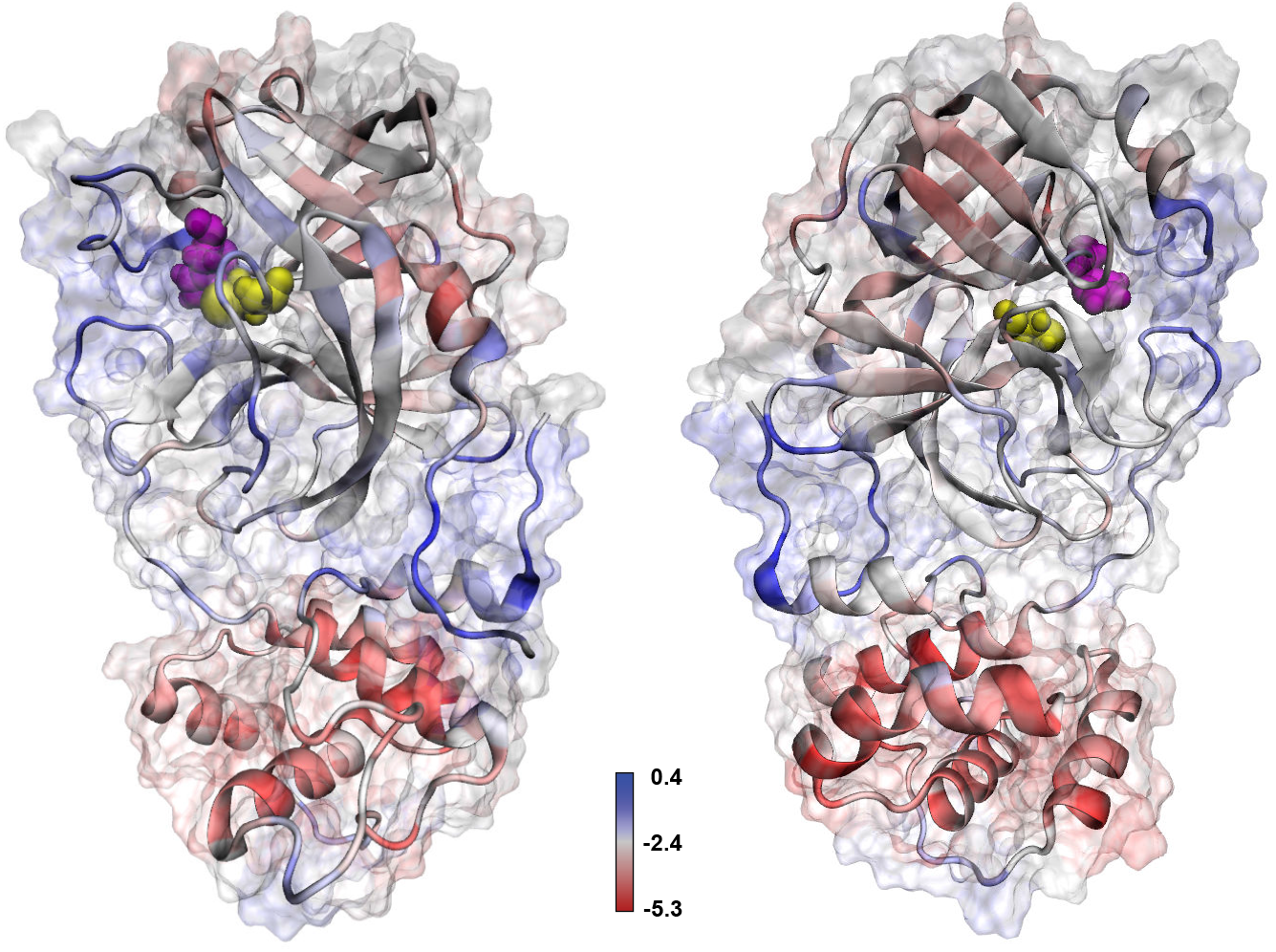
Local variation indices for M^*pro*^ backbone torsion angles (front/back views). Blue residues show higher levels of cross-variant *ϕ, Ψ* differences relative to baseline variation; red residues show little evidence of structural difference across variants. Domain 3 is substantially conserved, while greater change is seen in the inter-domain regions and loop regions adjacent to the active site.

The relatively high levels of conformational variation in the inter-domain regions suggest functionally relevant differences in global cohesion across variants. To assess this, we employ protein structure networks (PSNs), which are well-suited to assessing the looseness or cohesiveness of contacts among chemical groups. Moiety-level PSNs were constructed for each frame within each variant trajectory, using the definitions of (*32*) (Supplementary Figure S4). The assessment of global cohesion was performed by computing the mean degree *k*-core number for all moieties in each structure; to allow comparison of global cohesion within domains, we also compute mean core numbers within each of the three domains. The mean core number can be considered an index of structural cohesion, with higher values indicating greater numbers of redundant contacts among chemical groups (*33*). To account for within-trajectory autocorrelation in comparing mean core numbers, autocorrelation-corrected parametric bootstrap confidence intervals and standard errors were employed.

Figure 4 shows global and domain-specific cohesion levels (i.e., mean core numbers) for all variants, sorted in descending order of mean cohesion. (Means and standard errors for each variant can be found in Supplementary Table S1.) As suggested from the torsion angle analysis, cohesion differs significantly among variants, both globally and within domains. On average, the majority of variants are estimated to be less cohesively structured than WT, with the exception of domain 3 (in which WT does not differ significantly from the mean). It is possible that these differences indicate selection for more globally flexible structures (again, with the exception of domain 3). Whether or not this is the case, however, it appears clear that less cohesive structures are not strongly selected *against*. Such flexibility may affect dimerization kinetics, which is potentially relevant to the development of robust dimerization inhibitors.

**Figure 4:**
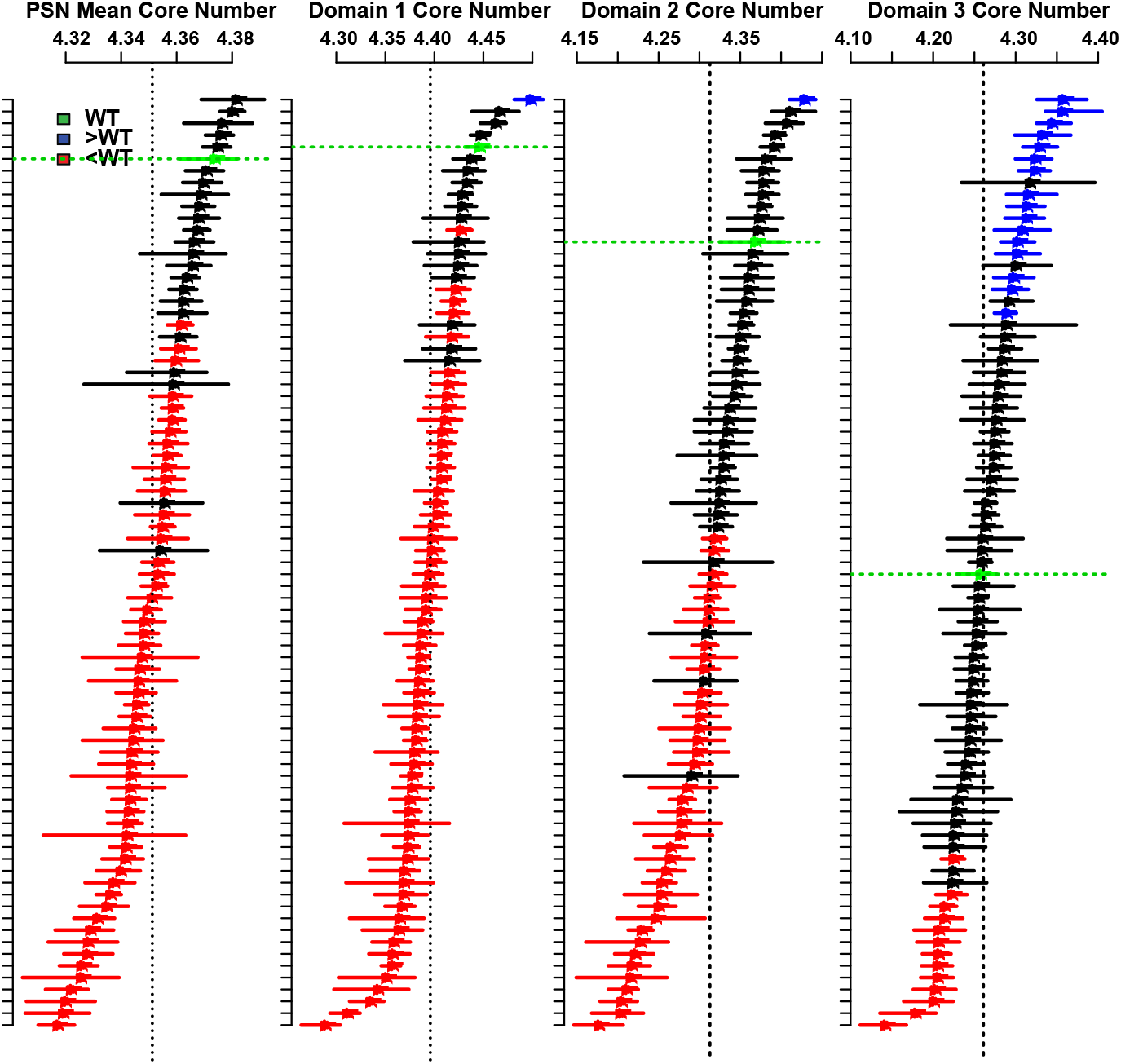
Mean core numbers for M^*pro*^ PSNs, by variant (ordering is by mean value in each panel). Points indicate trajectory means, with segments showing autocorrelation corrected 95% bootstrap confidence intervals; red/blue intervals have *t* values versus WT (green) of at least 2, indicating significant variation in structural cohesion across variants. Overall, the majority of variants are less cohesive than WT globally and in domains 1 and 2, while domain 3 cohesion in WT is typical of the variant set.

### Active site networks suggest potential activity differences across M^*pro*^ variants

The observation of structural variation in loop regions associated with the binding pocket motivates closer examination of variation in the M^*pro*^ active site. To this end, subgraphs of the full protein structure networks comprising moieties belonging to the active site residues and their neighbors were constructed to produce *active site networks* (ASNs) (*34*) for all conformations. A protein’s ASN describes physical interactions among active site moieties and other groups that are immediately adjacent in the 3D structure, irrespective of their positions in the amino acid sequence. Per (*34*), we compute for each ASN a *constraint score,* a general measure of active site flexibility that is associated with substrate specificity. The constraint score is the first principal component of a set of several network metrics (see Methods), with higher values indicating a greater tendency for the catalytic residues to be constrained by cohesive contacts with other residues, and lower values indicating fewer such constraints. Examples of ASNs corresponding to the maximum, minimum, and mean observed constraint values over all observed M^*pro*^ conformations are shown in Fig. 5.

**Figure 5:**
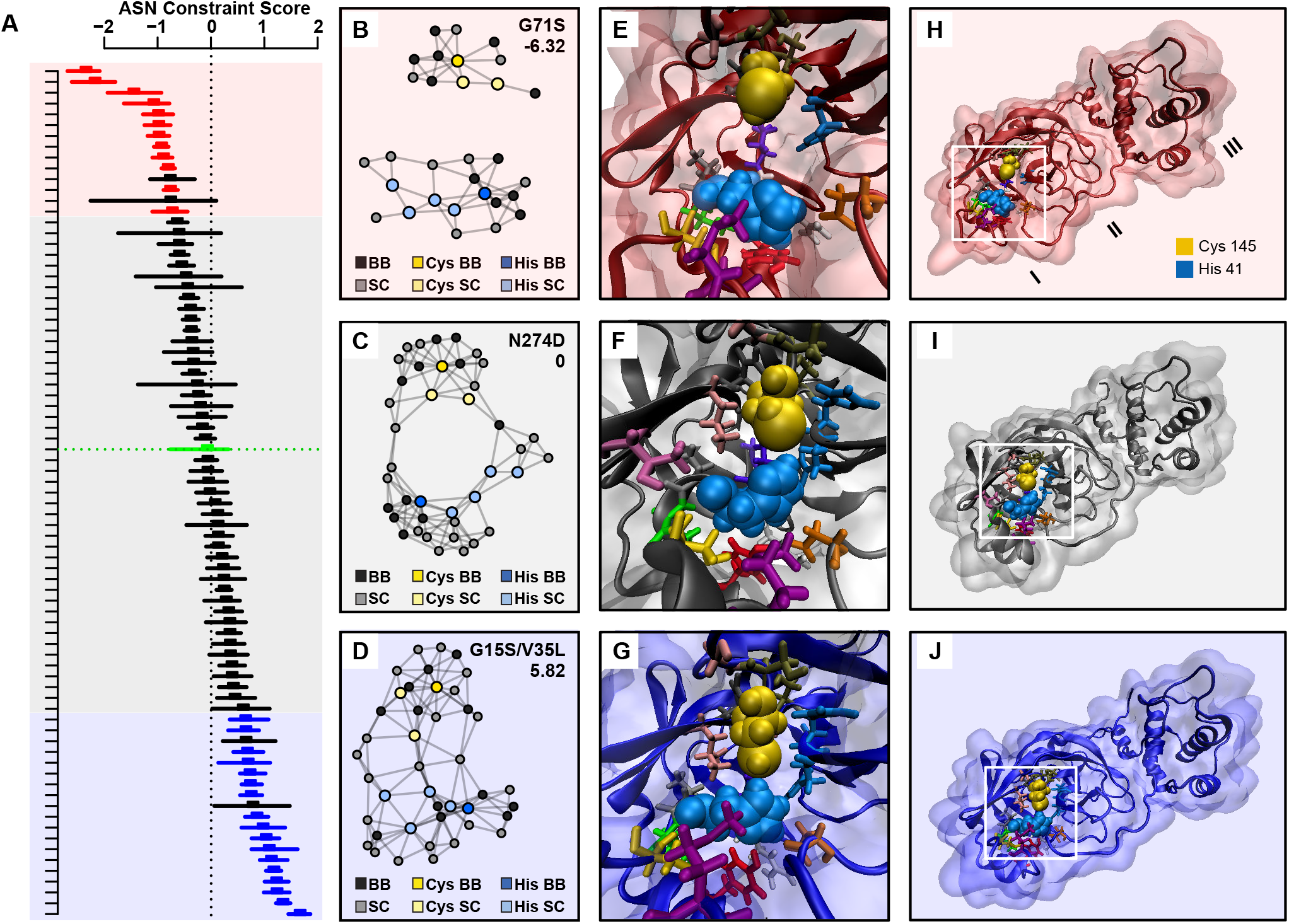
A. Mean active site constraint scores and 95% autocorrelation corrected parametric bootstrap confidence intervals, by variant. Higher values indicate greater constraints on active site residues; red/blue intervals have *t* values versus WT (green) of at least 2, indicating significant variation in average constraint across variants. B. Minimum, C. mean and D. maximum constraint ASNs over all frames. Low constraint conformations are characterized by no shared partners between the catalytic residues (colored nodes), while highly constrained conformations show cohesively reinforced contacts between them.

Examination of the mean constraint scores for each variant trajectory suggests potential activity differences across M^*pro*^ variants. Fig. 5A shows mean constraint scores for each variant, with autocorrelation-corrected parametric bootstrap confidence intervals. Of the 79 trajectories examined, 22 (28%) were significantly below the grand mean (dotted vertical line) and 28 (35%) were significantly above it; similarly, when directly compared to WT, 12 variants were observed to be significantly less constrained, while 17 were significantly more constrained (i.e., bootstrap *t*-scores less than −2 or greater than 2, respectively). 43 out of 78 variants (55%) showed nominally higher levels of mean constraint than WT (discounting significance), suggesting a lack of uniform selection pressure for active sites that are more or less constrained than wild type (the fraction greater does not differ significantly from random deviation, *p* = 0.16, exact binomial test). Thus, although we do not see evidence here of systematic selection for net changes in active site constraint, we do see evidence that variants differ from each other and from WT in their average active site properties. These differences should be considered in the design of inhibitors that are robust to mutational change in M^*pro*^ over time. In particular, it is clear that the population of extant M^*pro*^ variants already possesses some phenotypic diversity in active site flexibility, potentially facilitating its ability to evolve around some types of inhibitors.

## Methods

### Sequence analysis and clustering

SARS-CoV-2 genome sequences were found by searching the GISAID (https://www.gisaid.org/) (*13*) EpiCoV database on 3 May, 2020, using the host keyword “human” and a cutoff date of 29 April, 2020, yielding a total of 15,432 SARS-CoV-2 genomes. Genomes outside the range of ± 3% reference (RefSeq: NC 045512.2) length (29,006bp – 30,800bp inclusive) or ≥ 1% N content were removed, leaving 10,644 “high-quality” sequences. Open reading frames in these high-quality full genomes were compared with a reference Mpro nucleotide sequence (WT, RefSeq: NC 045512.2, loc: 10,055–10,972), to extract Mpro sequences of at least 80% similarity using a script written in Python v3.7.0 (*35*). Genomes with gaps or ambiguous nucleotides (e.g. N, S, D, per International Union of Pure and Applied Chemistry (IUPAC) nomenclature (*20*)), in the Mpro sequence were excluded from this data set, leaving a total of 10,578 sequences from high-coverage genomes.

Nucleotide sequences were converted into amino acid sequences and screened for nonsynonymous mutations against the WT M^*pro*^ using code written in Wolfram Mathematica 12.1 (*36*), yielding 511 non-synonymous mutations in M^*pro*^, 77 of which were unique. A single unique Mpro variant, found in a 24 April, 2020 dataset, but no longer available in the GISAID database, was also used in our analyses. Full genome alignments were performed using MUSCLE (*14*) on the complete set of non-synonymous Mpro mutants as well as reference WT, bat, and pangolin sequences. Trees were generated in MEGA X (*15*), using the Neighbor-Joining method (*37*); a bootstrap test (*38*) of 1000 replicates was performed, and distances were calculated using the Maximum Composite Likelihood model (*38*). In all, 515 full genomes were used in phylogenetic analyses; 78 unique Mpro mutants and a reference WT sequence (79 total) were used for molecular modeling.

### Molecular modeling of wild-type and variant protein structures

Initial conditions for the WT trajectory used here are based on the A monomer of PDB structure 6Y2E (*39*), representing a mature (i.e., cleaved pro-sequence) protein. Initial variant protein structures were predicted using MODELLER 9.23 (*30*), using the 6Y2E structure as a template; three rounds of annealing and MD refinement were performed using the “slow” optimization level for each. Initial structures were then processed to correct protonation states to reflect their predicted cellular environment (with protonation states predicted using PROPKA 3.1 (*40*)). Each corrected model structure was then minimized and equilibrated in explicit solvent; simulations were performed using NAMD (*41*) with the CHARMM36 forcefield (*42*) in TIP3P water (*43*) at 310 K under periodic boundary conditions (with a 10 Å margin water box). Solvated protein models were energy-minimized for 10,000 iterations before being simulated for 0.5ns to adjust water box size, after which a 10ns trajectory was simulated with conformations being sampled every 20ps; an *N_p_T* ensemble was used, with temperature controlled via Langevin dynamics with a damping coefficient of 1/ps and Nosé-Hoover Langevin piston pressure control set to 1 atm (*44, 45*).

### Network analysis

A protein structure network (PSN) was calculated for each modeled conformation of each variant via scripts employing the `statnet` (*46–48*), `Rpdb` (*49*), and `bio3d` (*50*) libraries for R (*51*). Vertices were defined using the method of (*32*), where each node represents a chemical moieity, with edges being defined by interatomic contacts. Specifically, two nodes *i* and *j* are considered adjacent if *i* contains atom *g* and *j* contains atom *h* such that the *g, h* distance is less than 1.1 times the sum of their respective van der Waals radii (using values from (*52*)). The node definitions are illustrated in Supplementary Figure S4A, and a small-moiety PSN of this type for WT M^*pro*^ is shown in Supplementary Figure S4B. Active site networks (ASNs) were constructed from each PSN as described in (*34*). Briefly, all vertices belonging to the catalytic Cys and His residues were identified, along with all vertices adjacent to these vertices within the PSN. The ASN was then defined as the subgraph of the corresponding PSN induced by this combined vertex set (Supplementary Figure S4C.)

To assess overall cohesion, degree *k*-core values (*53*) were calculated for each vertex in each PSN, and the average core number was computed for the entire protein and for the vertices in each domain, respectively. All calculations were performed using the `sna` library (*48*) for R. For each vertex associated with a moiety in the active site, three measures identified as associated with active site constraint by (*34*) were computed: the degree, or number of ties to other vertices; the triangle degree, or number of triangles (3-cliques) to which the vertex belongs; and core number, or number of the highest degree k-core (*54*) to which the vertex belongs. Physically, these respectively indicate the total number of contacts associated with the chemical group (potentially impeding its motion), the number of truss-like, triangular structures in which the group is embedded (again, restricting mobility), and the extent of local cohesion around the chemical group, which is found to distinguish “tighter” and “looser” packing regimes (*33*). To summarize the impact of each measure over the active site as a whole, values were averaged across active-site vertices. As an an additional constraint measure, the number of paths between each pair of active-site vertices through neighboring (i.e., non-active site) vertices was computed, and the log of the minimum of this value over the set of active site vertex pairs was employed as a measure of site cohesion. Intuitively, high values of site cohesion indicate that all active site chemical groups are connected by a large number of indirect contacts, while low values suggest that at least one pair of active site moieties has few local pathways holding them together. These four indices (mean active site degree, mean active site triangle degree, mean active site core number, and site cohesion) were used to produce an omnibus index of site constraint via principal component analysis (PCA) of the standardized network measures over all modeled conformations, per the approach of (*34*). This first principal component (the constraint score) accounted for approximately 71% of the variance in all four measures, and the ratio of its associated eigenvalue to the next largest was approximately 4.7 (confirming the dominance of the principal eigenvector).

#### Comparing mean cohesion and constraint scores across variants

Because cohesion and constraint scores are heavily autocorrelated within trajectories, we employ a parametric bootstrap strategy to obtain autocorrelation-corrected standard errors and confidence intervals (*55*). For each time series of scores for each trajectory, an autoregressive (AR) model with AIC-selected order was fit, and the estimated series mean obtained. (Estimation performed by maximum likelihood estimation using the `ar` function in R (*51*).) The whitened residuals from the time series model were then used to construct 5,000 parametric bootstrap replicate series, which were then re-fit to obtain bootstrap replicate means. Mean estimates from the bootstrap replicates were used to construct 95% bootstrap confidence intervals and standard errors for the series mean, as shown in Figs. 4 and 5. This procedure was applied to the MD trajectory for each variant. For the WT comparisons shown in Figs. 4 and 5, *t* values for mean constraint or cohesion score of each variant trajectory versus WT were constructed using the bootstrap-estimated standard errors, with variant trajectories indicated in red or blue (respectively) if the differences of their mean scores versus WT led to *t* statistics below −2 or above 2. For cohesion scores, mean and bootstrap standard errors are provided for the full protein and each domain in Supplementary Table S1.

## Author Contributions

R.W.M. and C.T.B. designed the study. T.J.C., G.R.T., M.G.C., and R.W.M analyzed sequence data. E.M.D. and C.T.B developed, implemented, and analyzed the simulation and network studies. M.G.C., V.F., S.Z., C.T.B., and R.W.M. performed structural analysis. T.J.C., G.R.T., C.T.B., and R.W.M. wrote the manuscript.

## Acknowledgments

This research was supported by NSF awards DMS-1361425 to C.T.B. and R.W.M. and IIS-1939237 to C.T.B., NASA Award 80NSSC20K0620 to R.W.M. and C.T.B., NIH grant 2R01EY021514 to RWM, the CIFAR Molecular Architecture of Life program, and the UC Irvine School of Physical Sciences Dean’s Excellence Fund. R.W.M. is a CIFAR Fellow.

## Supplementary materials

### Table of Contents

#### Figures

- Molecular models of representative variants (S1);
- Sites of known M^*pro*^ mutations, mapped on the wild-type structure (S2);
- Additional information on the fibril model reference measure (S3);
- Local variation index values for M^*pro*^ backbone torsion angle, by residue number (S4);
- Additional detail on the PSNs used in this work (S5)

#### Tables

- Mean cohesion scores (*k*-core number) and autocorrelation-corrected bootstrap standard errors by variant (S1);
- Accession numbers, locations, and dates of collection of all M^*pro*^ variants referred to in this work (S2)

#### Additional files available for download

- Uncompressed version of the tree depicted in Figure 1 (muscle gisaid 20200429.txt);
- Full acknowledgments for all sequences used in this work (514 acknowledgements.pdf)

---

**Figure S1:**
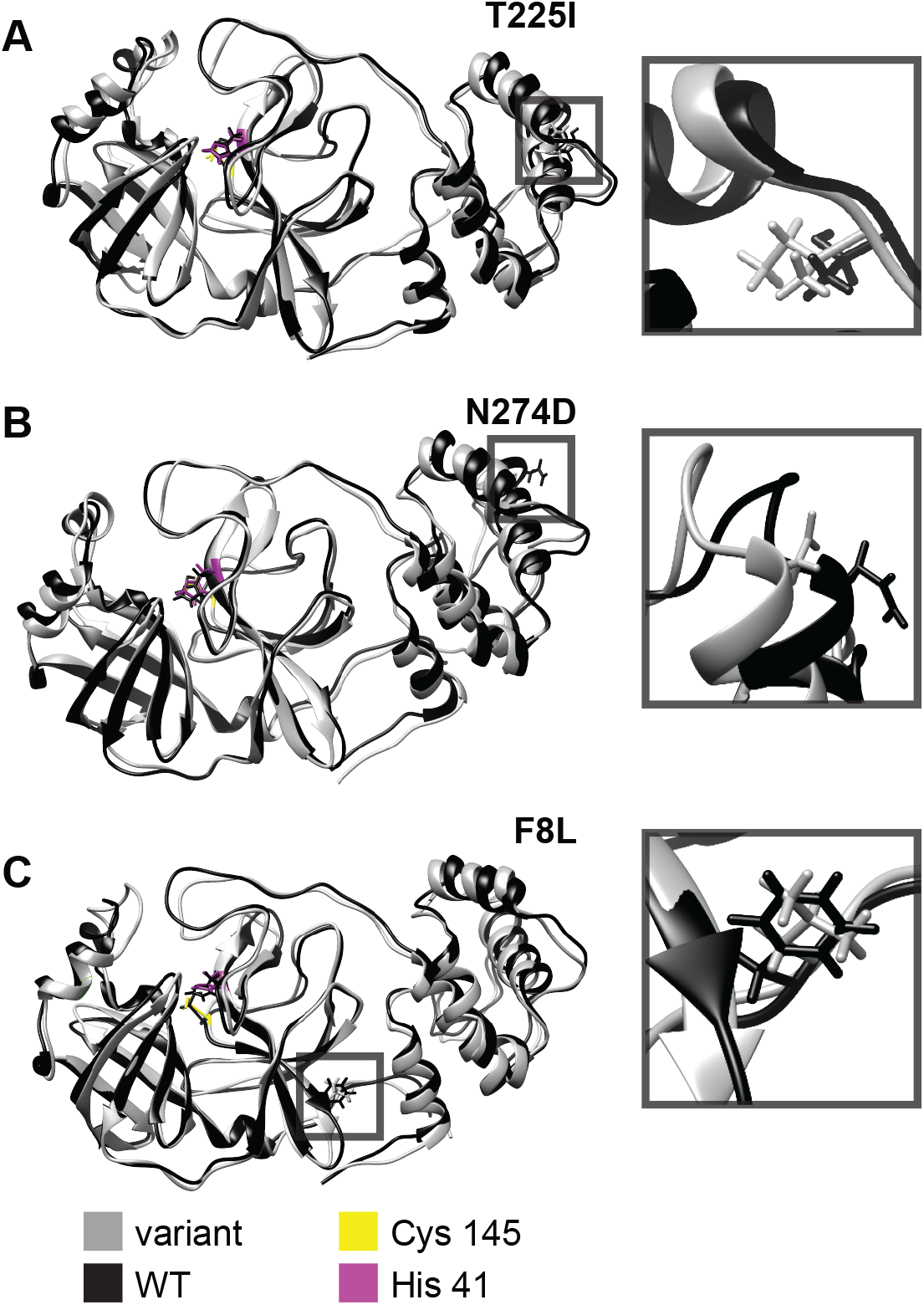
Molecular models of representative variants are shown in gray, overlaid with WT in black. The side chains are shown for active site residues and mutation sites. A. T225I B. N274D C. F8L.

**Figure S2:**
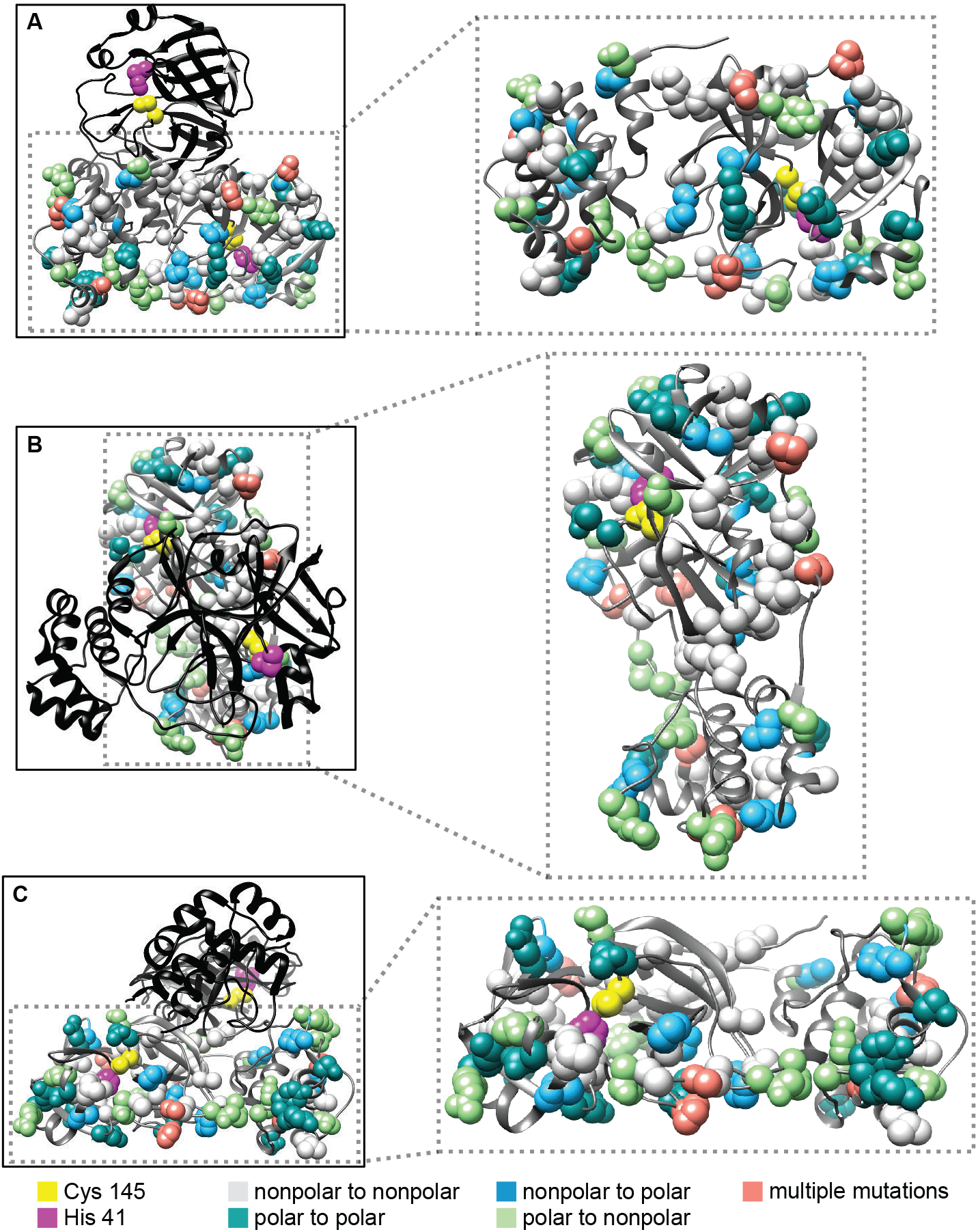
The positions of each mutated residue are shown plotted on the wild-type protein (PDB ID: (*31*). Panels A-C show different views of the M^*pro*^ dimer (left) and monomer (right.) One chain of the dimer is shown in black; on this monomer, only the active site residues His 41 (magenta) and Cy 145 (yellow) are shown in space-filling representations. On the section monomer (gray) side-chains of the mutated residues are also shown, using the following color coding to indicate the properties of the substituted residue: light gray - polar to nonpolar; teal - polar to polar; sky blue - nonpolar to polar; light green - polar to nonpolar; and salmon - multiple mutations (i.e. two or more independent substitutions with different properties.)

**Table S1:**
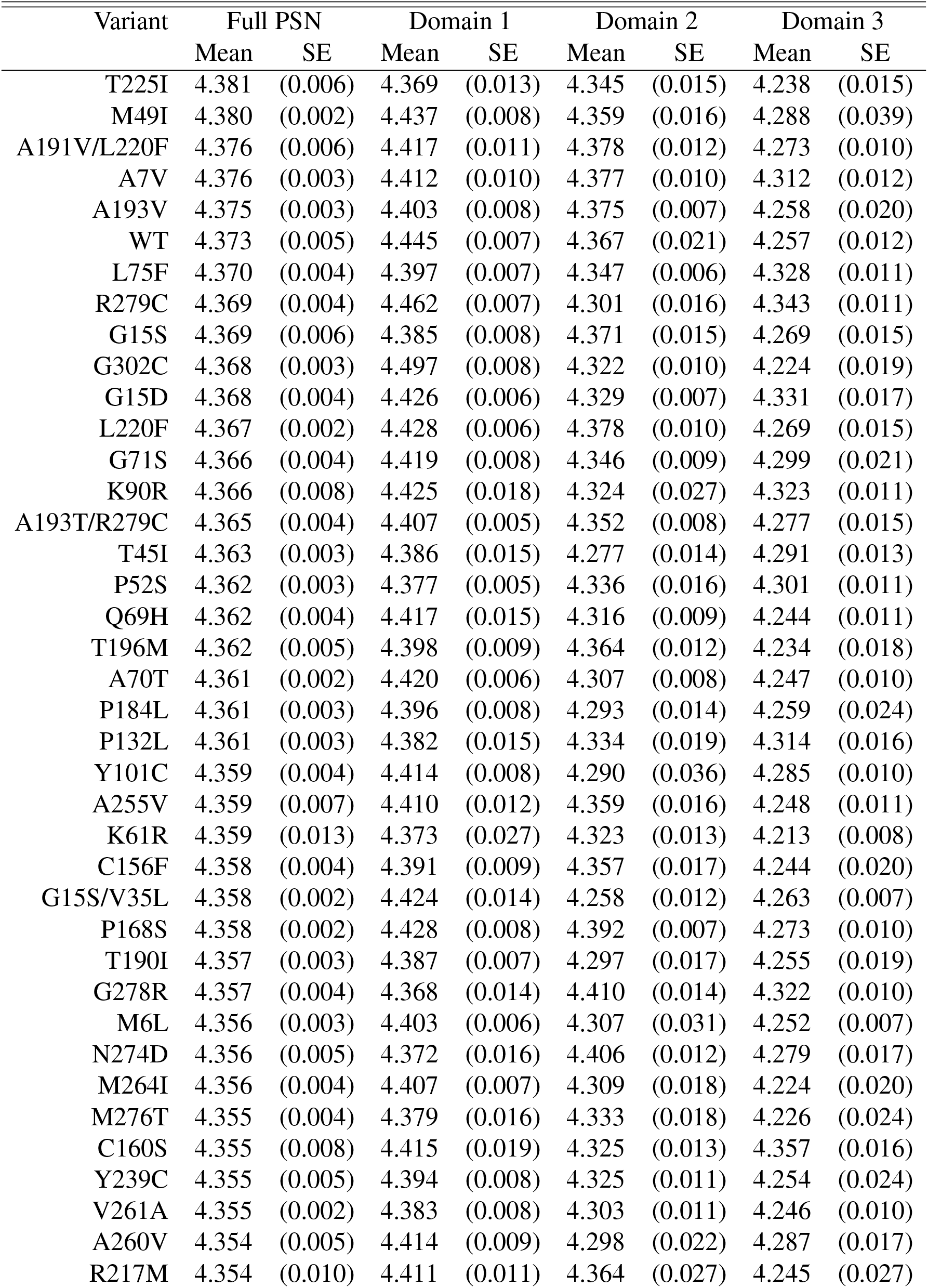
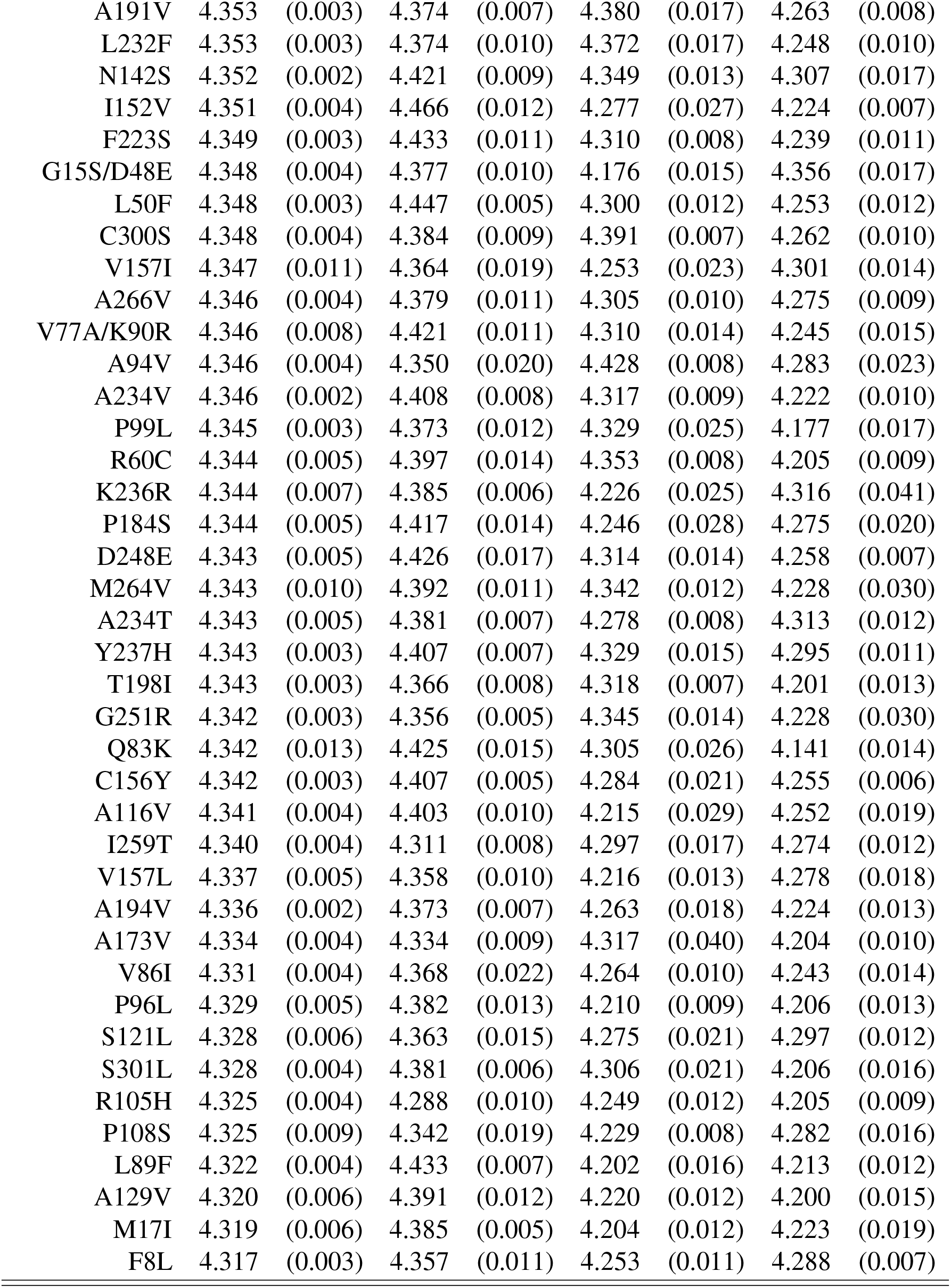
Mean cohesion scores (*k*-core number) and autocorrelation-corrected bootstrap standard errors by variant, for the entire PSN and by domain.

| Variant | Full PSN |  | Domain 1 |  | Domain 2 |  | Domain 3 |  |
| --- | --- | --- | --- | --- | --- | --- | --- | --- |
|  | Mean | SE | Mean | SE | Mean | SE | Mean | SE |
| T225I | 4.381 | (0.006) | 4.369 | (0.013) | 4.345 | (0.015) | 4.238 | (0.015) |
| M49I | 4.380 | (0.002) | 4.437 | (0.008) | 4.359 | (0.016) | 4.288 | (0.039) |
| A191V/L220F | 4.376 | (0.006) | 4.417 | (0.011) | 4.378 | (0.012) | 4.273 | (0.010) |
| A7V | 4.376 | (0.003) | 4.412 | (0.010) | 4.377 | (0.010) | 4.312 | (0.012) |
| A193V | 4.375 | (0.003) | 4.403 | (0.008) | 4.375 | (0.007) | 4.258 | (0.020) |
| WT | 4.373 | (0.005) | 4.445 | (0.007) | 4.367 | (0.021) | 4.257 | (0.012) |
| L75F | 4.370 | (0.004) | 4.397 | (0.007) | 4.347 | (0.006) | 4.328 | (0.011) |
| R279C | 4.369 | (0.004) | 4.462 | (0.007) | 4.301 | (0.016) | 4.343 | (0.011) |
| G15S | 4.369 | (0.006) | 4.385 | (0.008) | 4.371 | (0.015) | 4.269 | (0.015) |
| G302C | 4.368 | (0.003) | 4.497 | (0.008) | 4.322 | (0.010) | 4.224 | (0.019) |
| G15D | 4.368 | (0.004) | 4.426 | (0.006) | 4.329 | (0.007) | 4.331 | (0.017) |
| L220F | 4.367 | (0.002) | 4.428 | (0.006) | 4.378 | (0.010) | 4.269 | (0.015) |
| G71S | 4.366 | (0.004) | 4.419 | (0.008) | 4.346 | (0.009) | 4.299 | (0.021) |
| K90R | 4.366 | (0.008) | 4.425 | (0.018) | 4.324 | (0.027) | 4.323 | (0.011) |
| A193T/R279C | 4.365 | (0.004) | 4.407 | (0.005) | 4.352 | (0.008) | 4.277 | (0.015) |
| T45I | 4.363 | (0.003) | 4.386 | (0.015) | 4.277 | (0.014) | 4.291 | (0.013) |
| P52S | 4.362 | (0.003) | 4.377 | (0.005) | 4.336 | (0.016) | 4.301 | (0.011) |
| Q69H | 4.362 | (0.004) | 4.417 | (0.015) | 4.316 | (0.009) | 4.244 | (0.011) |
| T196M | 4.362 | (0.005) | 4.398 | (0.009) | 4.364 | (0.012) | 4.234 | (0.018) |
| A70T | 4.361 | (0.002) | 4.420 | (0.006) | 4.307 | (0.008) | 4.247 | (0.010) |
| P184L | 4.361 | (0.003) | 4.396 | (0.008) | 4.293 | (0.014) | 4.259 | (0.024) |
| P132L | 4.361 | (0.003) | 4.382 | (0.015) | 4.334 | (0.019) | 4.314 | (0.016) |
| Y101C | 4.359 | (0.004) | 4.414 | (0.008) | 4.290 | (0.036) | 4.285 | (0.010) |
| A255V | 4.359 | (0.007) | 4.410 | (0.012) | 4.359 | (0.016) | 4.248 | (0.011) |
| K61R | 4.359 | (0.013) | 4.373 | (0.027) | 4.323 | (0.013) | 4.213 | (0.008) |
| C156F | 4.358 | (0.004) | 4.391 | (0.009) | 4.357 | (0.017) | 4.244 | (0.020) |
| G15S/V35L | 4.358 | (0.002) | 4.424 | (0.014) | 4.258 | (0.012) | 4.263 | (0.007) |
| P168S | 4.358 | (0.002) | 4.428 | (0.008) | 4.392 | (0.007) | 4.273 | (0.010) |
| T190I | 4.357 | (0.003) | 4.387 | (0.007) | 4.297 | (0.017) | 4.255 | (0.019) |
| G278R | 4.357 | (0.004) | 4.368 | (0.014) | 4.410 | (0.014) | 4.322 | (0.010) |
| M6L | 4.356 | (0.003) | 4.403 | (0.006) | 4.307 | (0.031) | 4.252 | (0.007) |
| N274D | 4.356 | (0.005) | 4.372 | (0.016) | 4.406 | (0.012) | 4.279 | (0.017) |
| M264I | 4.356 | (0.004) | 4.407 | (0.007) | 4.309 | (0.018) | 4.224 | (0.020) |
| M276T | 4.355 | (0.004) | 4.379 | (0.016) | 4.333 | (0.018) | 4.226 | (0.024) |
| C160S | 4.355 | (0.008) | 4.415 | (0.019) | 4.325 | (0.013) | 4.357 | (0.016) |
| Y239C | 4.355 | (0.005) | 4.394 | (0.008) | 4.325 | (0.011) | 4.254 | (0.024) |
| V261A | 4.355 | (0.002) | 4.383 | (0.008) | 4.303 | (0.011) | 4.246 | (0.010) |
| A260V | 4.354 | (0.005) | 4.414 | (0.009) | 4.298 | (0.022) | 4.287 | (0.017) |
| R217M | 4.354 | (0.010) | 4.411 | (0.011) | 4.364 | (0.027) | 4.245 | (0.027) |
|  | Mean | SE | Mean | SE | Mean | SE | Mean | SE |
| A191V | 4.353 | (0.003) | 4.374 | (0.007) | 4.380 | (0.017) | 4.263 | (0.008) |
| L232F | 4.353 | (0.003) | 4.374 | (0.010) | 4.372 | (0.017) | 4.248 | (0.010) |
| N142S | 4.352 | (0.002) | 4.421 | (0.009) | 4.349 | (0.013) | 4.307 | (0.017) |
| I152V | 4.351 | (0.004) | 4.466 | (0.012) | 4.277 | (0.027) | 4.224 | (0.007) |
| F223S | 4.349 | (0.003) | 4.433 | (0.011) | 4.310 | (0.008) | 4.239 | (0.011) |
| G15S/D48E | 4.348 | (0.004) | 4.377 | (0.010) | 4.176 | (0.015) | 4.356 | (0.017) |
| L50F | 4.348 | (0.003) | 4.447 | (0.005) | 4.300 | (0.012) | 4.253 | (0.012) |
| C300S | 4.348 | (0.004) | 4.384 | (0.009) | 4.391 | (0.007) | 4.262 | (0.010) |
| V157I | 4.347 | (0.011) | 4.364 | (0.019) | 4.253 | (0.023) | 4.301 | (0.014) |
| A266V | 4.346 | (0.004) | 4.379 | (0.011) | 4.305 | (0.010) | 4.275 | (0.009) |
| V77A/K90R | 4.346 | (0.008) | 4.421 | (0.011) | 4.310 | (0.014) | 4.245 | (0.015) |
| A94V | 4.346 | (0.004) | 4.350 | (0.020) | 4.428 | (0.008) | 4.283 | (0.023) |
| A234V | 4.346 | (0.002) | 4.408 | (0.008) | 4.317 | (0.009) | 4.222 | (0.010) |
| P99L | 4.345 | (0.003) | 4.373 | (0.012) | 4.329 | (0.025) | 4.177 | (0.017) |
| R60C | 4.344 | (0.005) | 4.397 | (0.014) | 4.353 | (0.008) | 4.205 | (0.009) |
| K236R | 4.344 | (0.007) | 4.385 | (0.006) | 4.226 | (0.025) | 4.316 | (0.041) |
| P184S | 4.344 | (0.005) | 4.417 | (0.014) | 4.246 | (0.028) | 4.275 | (0.020) |
| D248E | 4.343 | (0.005) | 4.426 | (0.017) | 4.314 | (0.014) | 4.258 | (0.007) |
| M264V | 4.343 | (0.010) | 4.392 | (0.011) | 4.342 | (0.012) | 4.228 | (0.030) |
| A234T | 4.343 | (0.005) | 4.381 | (0.007) | 4.278 | (0.008) | 4.313 | (0.012) |
| Y237H | 4.343 | (0.003) | 4.407 | (0.007) | 4.329 | (0.015) | 4.295 | (0.011) |
| T198I | 4.343 | (0.003) | 4.366 | (0.008) | 4.318 | (0.007) | 4.201 | (0.013) |
| G251R | 4.342 | (0.003) | 4.356 | (0.005) | 4.345 | (0.014) | 4.228 | (0.030) |
| Q83K | 4.342 | (0.013) | 4.425 | (0.015) | 4.305 | (0.026) | 4.141 | (0.014) |
| C156Y | 4.342 | (0.003) | 4.407 | (0.005) | 4.284 | (0.021) | 4.255 | (0.006) |
| A116V | 4.341 | (0.004) | 4.403 | (0.010) | 4.215 | (0.029) | 4.252 | (0.019) |
| I259T | 4.340 | (0.004) | 4.311 | (0.008) | 4.297 | (0.017) | 4.274 | (0.012) |
| V157L | 4.337 | (0.005) | 4.358 | (0.010) | 4.216 | (0.013) | 4.278 | (0.018) |
| A194V | 4.336 | (0.002) | 4.373 | (0.007) | 4.263 | (0.018) | 4.224 | (0.013) |
| A173V | 4.334 | (0.004) | 4.334 | (0.009) | 4.317 | (0.040) | 4.204 | (0.010) |
| V86I | 4.331 | (0.004) | 4.368 | (0.022) | 4.264 | (0.010) | 4.243 | (0.014) |
| P96L | 4.329 | (0.005) | 4.382 | (0.013) | 4.210 | (0.009) | 4.206 | (0.013) |
| S121L | 4.328 | (0.006) | 4.363 | (0.015) | 4.275 | (0.021) | 4.297 | (0.012) |
| S301L | 4.328 | (0.004) | 4.381 | (0.006) | 4.306 | (0.021) | 4.206 | (0.016) |
| R105H | 4.325 | (0.004) | 4.288 | (0.010) | 4.249 | (0.012) | 4.205 | (0.009) |
| P108S | 4.325 | (0.009) | 4.342 | (0.019) | 4.229 | (0.008) | 4.282 | (0.016) |
| L89F | 4.322 | (0.004) | 4.433 | (0.007) | 4.202 | (0.016) | 4.213 | (0.012) |
| A129V | 4.320 | (0.006) | 4.391 | (0.012) | 4.220 | (0.012) | 4.200 | (0.015) |
| M17I | 4.319 | (0.006) | 4.385 | (0.005) | 4.204 | (0.012) | 4.223 | (0.019) |
|  | Mean | SE | Mean | SE | Mean | SE | Mean | SE |
| F8L | 4.317 | (0.003) | 4.357 | (0.011) | 4.253 | (0.011) | 4.288 | (0.007) |

**Figure S3:**
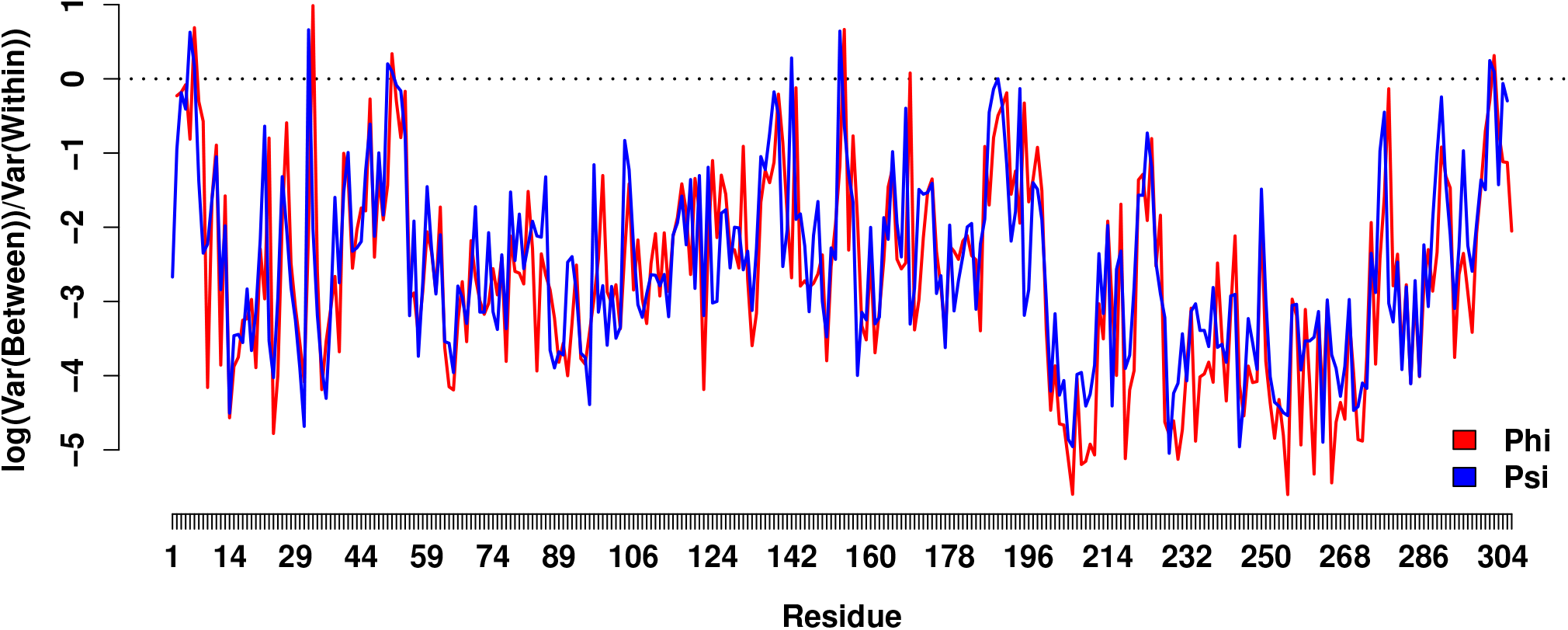
Local variation index values for M^*pro*^ backbone torsion angle, by residue number. While differences in mean angle between variants are generally smaller than angular variation within trajectories, some torsion angles show relatively high levels of variation net of dynamics; these are largely found in the interdomain interface, and loop regions near the active site.

**Figure S4:**
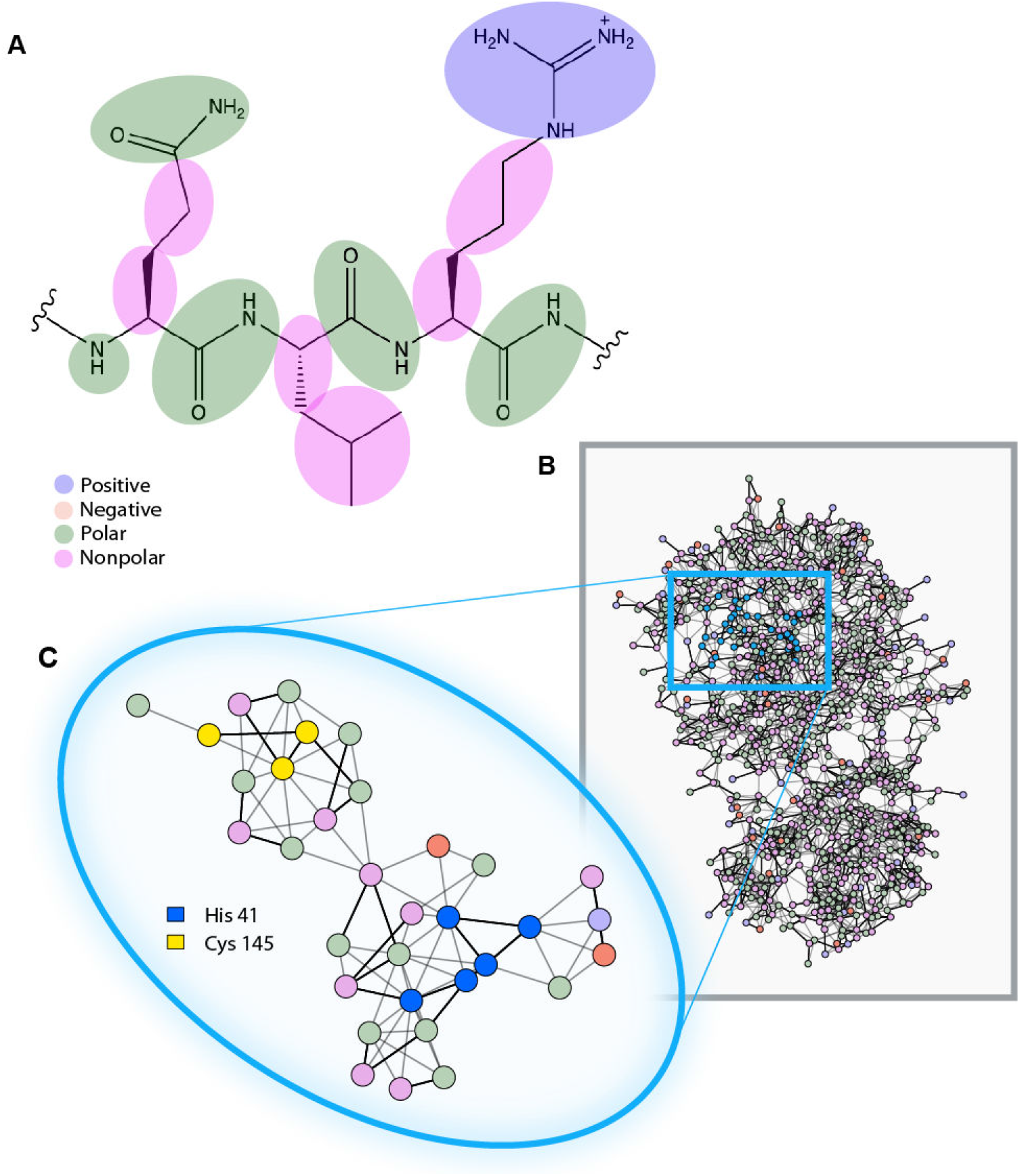
A. Node definitions of small-moiety networks defined as in (*32*), for the example of the peptide QLR. B. Small moiety PSN for WT M^*pro*^ (PDB ID: (*31*)), with nodes color-coded by chemical properties. C. ASN for WT M^*pro*^, which is a subgraph of the full PSN shown in Panel B.

**Table S2:** Accession numbers, locations, and dates of collection of variants as they appear in the uncompressed version of the tree depicted in Figure 1, which is available for download as a .txt file. Variants in bold are shown as individual branches in Figure 1. Those without accession numbers or dates represent subtrees that were compressed; their constituent variants reside underneath.

| Variant | Location | Accession | Date Collected |
| --- | --- | --- | --- |
| **Pangolin** | **Guangdong** | EPI_ISL_410721 | 2019 |
| **Bat** | **Yunnan** | EPI_ISL_402131 | 2013-07-24 |
| **K90R** | **Iceland** | EPI_ISL_424589 | 2020-03-28 |
| **K90R** | **Iceland** | EPI_ISL_424575 | 2020-03-28 |
| **V86I** | **Australia** |  |  |
| V86I | Australia | EPI_ISL_430478 | 2020-03-31 |
| V86I | Australia | EPI_ISL_426737 | 2020-03-26 |
| **K90R** | **Iceland** |  |  |
| K90R | Iceland | EPI_ISL_424489 | 2020-03-20 |
| K90R | Iceland | EPI_ISL_424407 | 2020-03-19 |
| K90R | Iceland | EPI_ISL_417623 | 2020-03-18 |
| K90R | Iceland | EPI_ISL_417613 | 2020-03-17 |
| K90R | Iceland | EPI_ISL_424498 | 2020-03-20 |
| K90R | Iceland | EPI_ISL_424475 | 2020-03-20 |
| K90R | Iceland | EPI_ISL_424406 | 2020-03-19 |
| K90R | Iceland | EPI_ISL_417650 | 2020-03-18 |
| K90R | Iceland | EPI_ISL_424500 | 2020-03-20 |
| K90R | Iceland | EPI_ISL_417600 | 2020-03-17 |
| K90R | Iceland | EPI_ISL_424393 | 2020-03-19 |
| K90R | Iceland | EPI_ISL_424447 | 2020-03-19 |
| K90R | Iceland | EPI_ISL_417625 | 2020-03-18 |
| K90R | Iceland | EPI_ISL_424508 | 2020-03-20 |
| K90R | Iceland | EPI_ISL_424511 | 2020-03-20 |
| K90R | Iceland | EPI_ISL_424380 | 2020-03-19 |
| K90R | Iceland | EPI_ISL_424392 | 2020-03-19 |
| K90R | Iceland | EPI_ISL_424419 | 2020-03-19 |
| K90R | Iceland | EPI_ISL_424460 | 2020-03-20 |
| K90R | Iceland | EPI_ISL_424512 | 2020-03-20 |
| K90R | Iceland | EPI_ISL_424515 | 2020-03-20 |
| K90R | Iceland | EPI_ISL_424516 | 2020-03-20 |
| K90R | Iceland | EPI_ISL_424568 | 2020-03-28 |
| K90R | Iceland | EPI_ISL_424601 | 2020-03-27 |
| K90R | Iceland | EPI_ISL_417790 | 2020-03-12 |
| K90R | Iceland | EPI_ISL_417836 | 2020-03-16 |
| K90R | Iceland | EPI_ISL_417536 | 2020-03-16 |
| K90R | Iceland | EPI_ISL_417549 | 2020-03-14 |
| K90R | Iceland | EPI_ISL_417612 | 2020-03-17 |
| K90R | Iceland | EPI_ISL_417630 | 2020-03-18 |
| K90R | Iceland | EPI_ISL_417635 | 2020-03-18 |
| K90R | Iceland | EPI_ISL_417636 | 2020-03-18 |
| K90R | Iceland | EPI_ISL_417645 | 2020-03-18 |
| K90R | Iceland | EPI_ISL_417678 | 2020-03-16 |
| K90R | Iceland | EPI_ISL_424414 | 2020-03-19 |
| K90R | Iceland | EPI_ISL_424450 | 2020-03-20 |
| K90R | Iceland | EPI_ISL_424461 | 2020-03-20 |
| K90R | Iceland | EPI_ISL_424483 | 2020-03-20 |
| K90R | Iceland | EPI_ISL_424573 | 2020-03-27 |
| K90R | Iceland | EPI_ISL_424576 | 2020-03-27 |
| K90R | Iceland | EPI_ISL_424605 | 2020-03-27 |
| K90R | Iceland | EPI_ISL_417786 | 2020-03-12 |
| K90R | Iceland | EPI_ISL_417703 | 2020-03-16 |
| K90R | Iceland | EPI_ISL_417808 | 2020-03-15 |
| K90R | Iceland | EPI_ISL_417809 | 2020-03-15 |
| K90R | Iceland | EPI_ISL_417827 | 2020-03-16 |
| K90R | Iceland | EPI_ISL_417543 | 2020-03-17 |
| K90R | Iceland | EPI_ISL_417573 | 2020-03-16 |
| K90R | Iceland | EPI_ISL_417583 | 2020-03-17 |
| K90R | Iceland | EPI_ISL_417588 | 2020-03-17 |
| K90R | Iceland | EPI_ISL_417626 | 2020-03-18 |
| K90R | Iceland | EPI_ISL_417634 | 2020-03-18 |
| K90R | Iceland | EPI_ISL_424449 | 2020-03-20 |
| K90R | Iceland | EPI_ISL_417755 | 2020-03-13 |
| K90R | Iceland | EPI_ISL_417712 | 2020-03-15 |
| K90R | Iceland | EPI_ISL_424593 | 2020-03-28 |
| K90R | Iceland | EPI_ISL_417709 | 2020-03-16 |
| K90R | Iceland | EPI_ISL_424440 | 2020-03-19 |
| K90R | Iceland | EPI_ISL_424582 | 2020-03-27 |
| K90R | Iceland | EPI_ISL_417753 | 2020-03-16 |
| K90R | Iceland | EPI_ISL_424559 | 2020-03-27 |
| K90R | Iceland | EPI_ISL_417680 | 2020-03-15 |
| K90R | Iceland | EPI_ISL_417810 | 2020-03-13 |
| K90R | Iceland | EPI_ISL_417633 | 2020-03-17 |
| K90R | Iceland | EPI_ISL_417745 | 2020-03-13 |
| K90R | Iceland | EPI_ISL_417597 | 2020-03-17 |
| K90R | Iceland | EPI_ISL_417741 | 2020-03-13 |
| K90R | Iceland | EPI_ISL_417676 | 2020-03-16 |
| K90R | Iceland | EPI_ISL_417739 | 2020-03-13 |
| K90R | Iceland | EPI_ISL_417586 | 2020-03-17 |
| K90R | Iceland | EPI_ISL_417542 | 2020-03-17 |
| K90R | Iceland | EPI_ISL_424619 | 2020-03-28 |
| K90R | Iceland | EPI_ISL_417774 | 2020-03-16 |
| K90R | Iceland | EPI_ISL_424474 | 2020-03-20 |
| K90R | Iceland | EPI_ISL_424584 | 2020-03-28 |
| K90R | Iceland | EPI_ISL_417824 | 2020-03-16 |
| K90R | Iceland | EPI_ISL_424571 | 2020-03-27 |
| K90R | Iceland | EPI_ISL_424588 | 2020-03-28 |
| K90R | Iceland | EPI_ISL_417552 | 2020-03-16 |
| K90R | Iceland | EPI_ISL_417537 | 2020-03-16 |
| **P184L** | **England** | EPI_ISL_420241 | 2020-03-26 |
| **R105H** | **Luxembourg** | EPI_ISL_419573 | 2020-03-11 |
| **P184S** | **England** | EPI_ISL_423288 | 2020-03-26 |
| **K90R** | **USA** | EPI_ISL_428779 | 2020-04-06 |
| **K90R** | **France** | EPI_ISL_420059 | 2020-03-21 |
| **P99L** | **Australia** | EPI_ISL_419756 | 2020-03-11 |
| **L232F** | **France** | EPI_ISL_421506 | 2020-03-21 |
| **K61R** | **USA** |  |  |
| K61R | USA | EPI_ISL_431014 | 2020-03-17 |
| K61R | USA | EPI_ISL_420306 | 2020-03-16 |
| K61R | USA | EPI_ISL_424346 | 2020-03-21 |
| K61R | USA | EPI_ISL_431016 | 2020-03-18 |
| K61R | USA | EPI_ISL_426459 | 2020-03-28 |
| K61R | USA | EPI_ISL_426464 | 2020-04-01 |
| K61R | USA | EPI_ISL_426502 | 2020-03-17 |
| **T196M** | **Iceland** |  |  |
| T196M | Iceland | EPI_ISL_417544 | 2020-03-17 |
| T196M | Iceland | EPI_ISL_424470 | 2020-03-20 |
| **T190I** | **Iceland** | EPI_ISL_424606 | 2020-03-28 |
| **A234T** | **USA** |  |  |
| A234T | USA | EPI_ISL_430327 | 2020-03-30 |
| A234T | USA | EPI_ISL_428777 | 2020-04-06 |
| A234T | USA | EPI_ISL_427484 | 2020-04-01 |
| **K90R** | **USA** | EPI_ISL_421301 | 2020-03-19 |
| **K90R** | **Scotland** | EPI_ISL_433228 | 2020-04-02 |
| **K90R** | **Wales** | EPI_ISL_432334 | 2020-04-04 |
| **K90R** | **England** | EPI_ISL_432988 | 2020-04-01 |
| **K90R** | **England** |  |  |
| K90R | England | EPI_ISL_423160 | 2020-03-24 |
| K90R | England | EPI_ISL_423161 | 2020-03-24 |
| K90R | England | EPI_ISL_421794 | 2020-03-26 |
| K90R | England | EPI_ISL_425436 | 2020-04-01 |
| K90R | England | EPI_ISL_421795 | 2020-03-26 |
| K90R | England | EPI_ISL_423152 | 2020-03-24 |
| **C300S** | **England** | EPI_ISL_433973 | 2020 |
| **G302C** | **England** | EPI_ISL_433732 | 2020-04-05 |
| **A193T/R279C** | **Australia** |  |  |
| A193T/R279C | Australia | EPI_ISL_426980 | 2020-03-28 |
| A193T/R279C | Australia | EPI_ISL_426989 | 2020-03-28 |
| A193T/R279C | Australia | EPI_ISL_426832 | 2020-03-25 |
| **R60C** | **England** |  |  |
| R60C | England | EPI_ISL_423509 | 2020-03-31 |
| R60C | England | EPI_ISL_420747 | 2020-03-28 |
| R60C | England | EPI_ISL_420748 | 2020-03-28 |
| **G15D** | **Belgium** | EPI_ISL_420422 | 2020-03-25 |
| **L89F** | **Georgia** | EPI_ISL_415643 | 2020-03-10 |
| **C160S** | **Turkey** | EPI_ISL_417413 | 2020-03-17 |
| **S301L** | **Australia** | EPI_ISL_427025 | 2020-03-31 |
| **K90R** | **Shanghai** |  |  |
| K90R | Shanghai | EPI_ISL_416332 | 2020-01-30 |
| K90R | Shanghai | EPI_ISL_416331 | 2020-01-30 |
| **P184S** | **Beijing** |  |  |
| P184S | Beijing | EPI_ISL_430734 | 2020-01-24 |
| P184S | Beijing | EPI_ISL_430736 | 2020-01-29 |
| P184S | Beijing | EPI_ISL_430735 | 2020-01-24 |
| P184S | Beijing | EPI_ISL_430742 | 2020-01-29 |
| **P184S** | **Malaysia** |  |  |
| P184S | Malaysia | EPI_ISL_430444 | 2020-02-12 |
| P184S | Malaysia | EPI_ISL_416907 | 2020-02-20 |
| **P184S** | **Beijing** | EPI_ISL_430729 | 2020-02-07 |
| **R105H** | **Australia** | EPI_ISL_419984 | 2020-03-23 |
| **V157I** | **Netherlands** | EPI_ISL_415503 | 2020-03-11 |
| **K90R** | **Australia** | EPI_ISL_426647 | 2020-03-20 |
| **Y237H** | **USA** | EPI_ISL_416720 | 2020-03-13 |
| **K90R** | **USA** | EPI_ISL_418873 | 2020-03-16 |
| **M264V** | **USA** | EPI_ISL_434304 | 2020-03-25 |
| **I259T** | **USA** | EPI_ISL_434193 | 2020-03-28 |
| **A194V** | **USA** | EPI_ISL_434184 | 2020-03-28 |
| **V157L** | **USA** | EPI_ISL_424196 | 2020-03-20 |
| **M276T** | **USA** | EPI_ISL_434109 | 2020-04-02 |
| **A255V** | **USA** |  |  |
| A255V | USA | EPI_ISL_424252 | 2020-03-23 |
| A255V | USA | EPI_ISL_418075 | 2020-03-15 |
| A255V | USA | EPI_ISL_424275 | 2020-03-22 |
| A255V | USA | EPI_ISL_424232 | 2020-03-21 |
| **A193V** | **USA** |  |  |
| A193V | USA | EPI_ISL_415610 | 2020-03-08 |
| A193V | USA | EPI_ISL_417350 | 2020-03-12 |
| **A173V** | **USA** | EPI_ISL_418082 | 2020-03-15 |
| **M264V** | **USA** | EPI_ISL_424297 | 2020-03-23 |
| **M264V** | **USA** | EPI_ISL_430924 | 2020-04-07 |
| **T190I** | **USA** | EPI_ISL_423007 | 2020-03-19 |
| **WT** | **Wuhan** | NC_045512 | 2019-12 |
| **A193V** | **USA** | EPI_ISL_429007 | 2020-03-18 |
| **P108S** | **Iceland** | EPI_ISL_424455 | 2020-03-20 |
| **R60C** | **Vietnam** |  |  |
| R60C | Vietnam | EPI_ISL_418269 | 2020-01-22 |
| R60C | Vietnam | EPI_ISL_418267 | 2020-01-22 |
| **I152V** | **Fujian** |  |  |
| I152V | Fujian | EPI_ISL_431784 | 2020-03-18 |
| I152V | Fujian | EPI_ISL_431783 | 2020-03-19 |
| **I259T** | **Netherlands** | EPI_ISL_422919 | 2020-03-19 |
| **V261A** | **England** | EPI_ISL_425498 | 2020-03-19 |
| **A7V** | **England** | EPI_ISL_425319 | 2020-03-28 |
| **Y239C** | **Scotland** |  |  |
| Y239C | Scotland | EPI_ISL_433339 | 2020-03-26 |
| Y239C | Scotland | EPI_ISL_433359 | 2020-03-27 |
| Y239C | Scotland | EPI_ISL_433357 | 2020-03-27 |
| Y239C | Scotland | EPI_ISL_433330 | 2020-03-26 |
| Y239C | Scotland | EPI_ISL_433341 | 2020-03-26 |
| Y239C | Scotland | EPI_ISL_433358 | 2020-03-27 |
| Y239C | Scotland | EPI_ISL_433307 | 2020-03-24 |
| **A191V** | **USA** | EPI_ISL_428258 | 2020-03-29 |
| **M6L** | **USA** |  |  |
| M6L | USA | EPI_ISL_428333 | 2020-03-27 |
| M6L | USA | EPI_ISL_428287 | 2020-03-23 |
| **V77A/K90R** | **England** | EPI_ISL_420520 | 2020-03-24 |
| **K90R** | **England** | EPI_ISL_420488 | 2020-03-20 |
| **K90R** | **Wales** | EPI_ISL_420937 | 2020-03-21 |
| **A116V** | **Wales** |  |  |
| A116V | Wales | EPI_ISL_432176 | 2020-04-01 |
| A116V | Wales | EPI_ISL_432248 | 2020-04-05 |
| **S301L** | **Wales** |  |  |
| S301L | Wales | EPI_ISL_422184 | 2020-03-27 |
| S301L | Wales | EPI_ISL_422052 | 2020-03-27 |
| **A260V** | **Australia** | EPI_ISL_427779 | 2020-03-24 |
| **F223S** | **Sweden** | EPI_ISL_429159 | 2020-03-10 |
| **G278R** | **England** | EPI_ISL_424003 | 2020-03-20 |
| **Q83K** | **USA** | EPI_ISL_430051 | 2020-04-02 |
| **T198I** | **Spain** | EPI_ISL_421515 | 2020-03-07 |
| **T225I** | **France** | EPI_ISL_414624 | 2020-02-26 |
| **Y101C** | **Germany** | EPI_ISL_425132 | 2020-03-23 |
| **D248E** | **Scotland** |  |  |
| D248E | Scotland | EPI_ISL_433545 | 2020-03-31 |
| D248E | Scotland | EPI_ISL_433543 | 2020-03-31 |
| D248E | Scotland | EPI_ISL_433593 | 2020-04-01 |
| D248E | Scotland | EPI_ISL_433534 | 2020-03-30 |
| D248E | Scotland | EPI_ISL_433366 | 2020-03-28 |
| D248E | Scotland | EPI_ISL_433546 | 2020-03-31 |
| D248E | Scotland | EPI_ISL_433542 | 2020-03-31 |
| D248E | Scotland | EPI_ISL_433569 | 2020-03-31 |
| D248E | Scotland | EPI_ISL_433139 | 2020-04-16 |
| D248E | Scotland | EPI_ISL_433241 | 2020-04-03 |
| D248E | Scotland | EPI_ISL_425982 | 2020-03-29 |
| D248E | Scotland | EPI_ISL_433113 | 2020-04-15 |
| D248E | Scotland | EPI_ISL_425794 | 2020-03-25 |
| D248E | Scotland | EPI_ISL_433603 | 2020-04-03 |
| D248E | Scotland | EPI_ISL_433400 | 2020-04-06 |
| D248E | Scotland | EPI_ISL_433509 | 2020-03-29 |
| D248E | Scotland | EPI_ISL_433505 | 2020-03-29 |
| D248E | Scotland | EPI_ISL_433502 | 2020-03-28 |
| D248E | Scotland | EPI_ISL_433398 | 2020-04-08 |
| D248E | Scotland | EPI_ISL_433333 | 2020-03-26 |
| D248E | Scotland | EPI_ISL_433653 | 2020-04-05 |
| D248E | Scotland | EPI_ISL_433598 | 2020-04-02 |
| D248E | Scotland | EPI_ISL_433641 | 2020-04-06 |
| D248E | Scotland | EPI_ISL_433632 | 2020-04-04 |
| D248E | Scotland | EPI_ISL_433626 | 2020-04-04 |
| D248E | Scotland | EPI_ISL_433302 | 2020-03-23 |
| D248E | Scotland | EPI_ISL_433126 | 2020-04-15 |
| D248E | Scotland | EPI_ISL_433355 | 2020-03-27 |
| D248E | Scotland | EPI_ISL_433617 | 2020-04-03 |
| **G15S/D48E** | **England** | EPI_ISL_425242 | 2020-03-31 |
| **G15S** | **England** | EPI_ISL_425300 | 2020-03-28 |
| **G15S** | **England** | EPI_ISL_433856 | 2020-04-08 |
| **G15S** | **England** | EPI_ISL_425443 | 2020-04-01 |
| **G15S** | **England** | EPI_ISL_423472 | 2020-03-31 |
| **G15S** | **Australia** |  |  |
| G15S | Australia | EPI_ISL_426662 | 2020-03-21 |
| G15S | Australia | EPI_ISL_427080 | 2020-03-23 |
| **G15S** | **Iceland** |  |  |
| G15S | Iceland | EPI_ISL_417581 | 2020-03-17 |
| G15S | Iceland | EPI_ISL_424510 | 2020-03-20 |
| G15S | Iceland | EPI_ISL_417643 | 2020-03-18 |
| G15S | Iceland | EPI_ISL_417642 | 2020-03-18 |
| **G15S** | **Costa Rica** | EPI_ISL_434538 | 2020-03-20 |
| **G15S** | **Denmark** | EPI_ISL_416140 | 2020-03-02 |
| **G15S** | **Wales** | EPI_ISL_421003 | 2020-03-23 |
| **G15S** | **Australia** | EPI_ISL_426914 | 2020-03-29 |
| **G15S** | **DRC** | EPI_ISL_420849 | 2020-03-28 |
| **G15S** | **England** | EPI_ISL_423071 | 2020-03-23 |
| **G15S** | **Wales** | EPI_ISL_415656 | 2020-03-12 |
| **G15S** | **England** | EPI_ISL_417265 | 2020-03-08 |
| **G15S** | **England** | EPI_ISL_423104 | 2020-03-23 |
| **G15S** | **England** | EPI_ISL_417307 | 2020-03-08 |
| **G15S** | **Finland** | EPI_ISL_418389 | 2020-03-13 |
| **G15S** | **England** | EPI_ISL_424121 | 2020-03-21 |
| **G15S** | **Sweden** | EPI_ISL_429116 | 2020-03-15 |
| **G15S** | **USA** | EPI_ISL_426626 | 2020-04-01 |
| **G15S** | **Wales** | EPI_ISL_421000 | 2020-03-23 |
| **G15S** | **Wales** | EPI_ISL_422186 | 2020-03-29 |
| **G15S** | **Wales** | EPI_ISL_413556 | 2020-03-04 |
| **G15S** | **England** |  |  |
| G15S | England | EPI_ISL_432502 | 2020-03-25 |
| G15S | England | EPI_ISL_420217 | 2020-03-27 |
| **G15S** | **Russia** | EPI_ISL_428881 | 2020-03-16 |
| **G15S** | **Argentina** | EPI_ISL_430809 | 2020-04-04 |
| **G15S** | **Argentina** | EPI_ISL_430808 | 2020-04-03 |
| **G15S** | **Russia** |  |  |
| G15S | Russia | EPI_ISL_428910 | 2020-03-31 |
| G15S | Russia | EPI_ISL_421275 | 2020-03-18 |
| **G15S** | **England** |  |  |
| G15S | England | EPI_ISL_432541 | 2020-04-03 |
| G15S | England | EPI_ISL_420215 | 2020-03-27 |
| G15S | England | EPI_ISL_420262 | 2020-03-24 |
| **G15S** | **Netherlands** | EPI_ISL_422682 | 2020-03-19 |
| **G15S** | **Netherlands** | EPI_ISL_422748 | 2020-03-25 |
| **G15S** | **England** | EPI_ISL_425248 | 2020-03-30 |
| **G15S** | **England** | EPI_ISL_433868 | 2020-04-08 |
| **G15S** | **England** | EPI_ISL_420723 | 2020-03-28 |
| **G15S** | **England** | EPI_ISL_433710 | 2020-04-04 |
| **G15S/V35L** | **Argentina** | EPI_ISL_430811 | 2020-04-11 |
| **G15S** | **Wales** | EPI_ISL_432302 | 2020-03-31 |
| **G15S** | **Denmark** | EPI_ISL_429273 | 2020-03-10 |
| **G15S** | **Australia** | EPI_ISL_427739 | 2020-03-21 |
| **G15S** | **Wales** |  |  |
| G15S | Wales | EPI_ISL_432295 | 2020-04-08 |
| G15S | Wales | EPI_ISL_432392 | 2020-04-08 |
| **G15S** | **Russia** |  |  |
| G15S | Russia | EPI_ISL_428889 | 2020-03-22 |
| G15S | Russia | EPI_ISL_428887 | 2020-03-22 |
| G15S | Russia | EPI_ISL_428874 | 2020-03-20 |
| G15S | Russia | EPI_ISL_428891 | 2020-03-25 |
| **G15S** | **Scotland** | EPI_ISL_433507 | 2020-03-29 |
| **G15S** | **Wales** | EPI_ISL_432273 | 2020-04-08 |
| **G15S** | **Iceland** | EPI_ISL_417538 | 2020-03-17 |
| **G15S** | **England** | EPI_ISL_423177 | 2020-03-24 |
| **G15S** | **England** | EPI_ISL_423495 | 2020-03-30 |
| **G15S** | **Scotland** |  |  |
| G15S | Scotland | EPI_ISL_425904 | 2020-03-22 |
| G15S | Scotland | EPI_ISL_425915 | 2020-03-23 |
| **G15S** | **Wales** | EPI_ISL_422094 | 2020-03-25 |
| **G15S** | **Wales** | EPI_ISL_422167 | 2020-03-27 |
| **G15S** | **Wales** | EPI_ISL_432378 | 2020-04-08 |
| **G15S** | **Wales** |  |  |
| G15S | Wales | EPI_ISL_432241 | 2020-04-08 |
| G15S | Wales | EPI_ISL_432345 | 2020-04-06 |
| G15S | Wales | EPI_ISL_432251 | 2020-04-04 |
| **G15S** | **Wales** | EPI_ISL_420977 | 2020-03-22 |
| **G15S** | **Wales** | EPI_ISL_432222 | 2020-03-30 |
| **G15S** | **England** | EPI_ISL_423175 | 2020-03-24 |
| **G15S** | **England** |  |  |
| G15S | England | EPI_ISL_421831 | 2020-03-23 |
| G15S | England | EPI_ISL_421824 | 2020-03-25 |
| G15S | England | EPI_ISL_421825 | 2020-03-25 |
| **G15S** | **England** | EPI_ISL_432549 | 2020-03-28 |
| **G15S** | **England** | EPI_ISL_432967 | 2020-03-31 |
| **G15S** | **England** |  |  |
| G15S | England | EPI_ISL_432691 | 2020-04-07 |
| G15S | England | EPI_ISL_432706 | 2020-04-08 |
| **G15S** | **England** | EPI_ISL_420199 | 2020-03-28 |
| **G15S** | **England** | EPI_ISL_420213 | 2020-03-29 |
| **G15S** | **England** | EPI_ISL_433798 | 2020-04-07 |
| **G15S** | **Scotland** | EPI_ISL_425761 | 2020-03-13 |
| **G15S** | **England** |  |  |
| G15S | England | EPI_ISL_420181 | 2020-03-27 |
| G15S | England | EPI_ISL_432837 | 2020-03-29 |
| G15S | England | EPI_ISL_420166 | 2020-03-25 |
| G15S | England | EPI_ISL_420244 | 2020-03-26 |
| G15S | England | EPI_ISL_420240 | 2020-03-23 |
| G15S | England | EPI_ISL_420278 | 2020-03-21 |
| G15S | England | EPI_ISL_420264 | 2020-03-23 |
| G15S | England | EPI_ISL_420261 | 2020-03-25 |
| G15S | England | EPI_ISL_432804 | 2020-03-27 |
| G15S | England | EPI_ISL_432712 | 2020-03-25 |
| **G15S** | **Scotland** |  |  |
| G15S | Scotland | EPI_ISL_433397 | 2020-04-08 |
| G15S | Scotland | EPI_ISL_433069 | 2020-04-13 |
| **G15S** | **England** | EPI_ISL_433493 | 2020-04-16 |
| **G15S** | **Australia** | EPI_ISL_427159 | 2020-03-30 |
| **G15S** | **England** | EPI_ISL_424119 | 2020-03-21 |
| **G15S** | **England** | EPI_ISL_424126 | 2020-03-21 |
| **G15S** | **Wales** | EPI_ISL_422180 | 2020-03-25 |
| **G15S** | **England** | EPI_ISL_423219 | 2020-03-25 |
| **G15S** | **Iceland** | EPI_ISL_417585 | 2020-03-17 |
| **G15S** | **Wales** |  |  |
| G15S | Wales | EPI_ISL_432230 | 2020-04-04 |
| G15S | Wales | EPI_ISL_432216 | 2020-04-06 |
| G15S | Wales | EPI_ISL_432237 | 2020-04-03 |
| **G15S** | **England** | EPI_ISL_424054 | 2020-03-09 |
| **G15S** | **England** | EPI_ISL_418770 | 2020-03-18 |
| **G15S** | **England** | EPI_ISL_423100 | 2020-03-24 |
| **G15S** | **Iceland** |  |  |
| G15S | Iceland | EPI_ISL_417783 | 2020-03-11 |
| G15S | Iceland | EPI_ISL_417539 | 2020-03-16 |
| **P108S** | **Iceland** | EPI_ISL_424466 | 2020-03-20 |
| **P52S** | **Russia** |  |  |
| P52S | Russia | EPI_ISL_428864 | 2020-03-11 |
| P52S | Russia | EPI_ISL_428863 | 2020-03-11 |
| **F8L** | **England** | EPI_ISL_433865 | 2020-04-08 |
| **R217M** | **England** | EPI_ISL_423428 | 2020-03-30 |
| **M17I** | **England** |  |  |
| M17I | England | EPI_ISL_423772 | 2020-03-22 |
| M17I | England | EPI_ISL_421903 | 2020-03-26 |
| **G71S** | **Denmark** | EPI_ISL_429484 | 2020-03-24 |
| **G71S** | **USA** | EPI_ISL_421352 | 2020-03-14 |
| **G71S** | **Germany** | EPI_ISL_412912 | 2020-02-25 |
| **G71S** | **Brazil** | EPI_ISL_416034 | 2020-03-04 |
| **G71S** | **Australia** | EPI_ISL_419893 | 2020-03-19 |
| **G71S** | **USA** | EPI_ISL_421621 | 2020-03-20 |
| **G71S** | **USA** | EPI_ISL_421351 | 2020-03-11 |
| **G71S** | **Australia** | EPI_ISL_427026 | 2020-03-30 |
| **G71S** | **USA** | EPI_ISL_421353 | 2020-03-14 |
| **G71S** | **USA** | EPI_ISL_421712 | 2020-03-18 |
| **G71S** | **USA** | EPI_ISL_421724 | 2020-03-18 |
| **G71S** | **Switzerland** | EPI_ISL_413021 | 2020-02-29 |
| **G251R** | **USA** | EPI_ISL_428732 | 2020-04-06 |
| **A70T** | **England** |  |  |
| A70T | England | EPI_ISL_433765 | 2020-04-05 |
| A70T | England | EPI_ISL_434038 | 2020-04-14 |
| **A234V** | **England** | EPI_ISL_425235 | 2020-03-30 |
| **A234V** | **England** | EPI_ISL_433681 | 2020-04-02 |
| **A234V** | **Australia** | EPI_ISL_427096 | 2020-04-03 |
| **M49I** | **Scotland** | EPI_ISL_425839 | 2020-03-13 |
| **V157L** | **Scotland** |  |  |
| V157L | Scotland | EPI_ISL_433428 | 2020-04-08 |
| V157L | Scotland | EPI_ISL_433256 | 2020-04-04 |
| **A129V** | **England** | EPI_ISL_433066 | 2020-04-05 |
| **L220F** | **Scotland** | EPI_ISL_433194 | 2020-04-19 |
| **A129V** | **Netherlands** | EPI_ISL_422860 | 2020-03-20 |
| **N274D** | **Scotland** |  |  |
| N274D | Scotland | EPI_ISL_433145 | 2020-04-16 |
| N274D | Scotland | EPI_ISL_433110 | 2020-04-15 |
| N274D | Scotland | EPI_ISL_433084 | 2020-04-12 |
| N274D | Scotland | EPI_ISL_433165 | 2020-04-18 |
| N274D | Scotland | EPI_ISL_433136 | 2020-04-16 |
| N274D | Scotland | EPI_ISL_433167 | 2020-04-18 |
| N274D | Scotland | EPI_ISL_433093 | 2020-04-14 |
| **C156Y** | **England** |  |  |
| C156Y | England | EPI_ISL_432938 | 2020-03-28 |
| C156Y | England | EPI_ISL_432935 | 2020-03-28 |
| **A94V** | **USA** |  |  |
| A94V | USA | EPI_ISL_422560 | 2020-03-22 |
| A94V | USA | EPI_ISL_427632 | 2020-04-06 |
| **S121L** | **USA** | EPI_ISL_430950 | 2020-03-29 |
| **L220F** | **USA** |  |  |
| L220F | USA | EPI_ISL_429987 | 2020-04-02 |
| L220F | USA | EPI_ISL_419256 | 2020-03-16 |
| L220F | USA | EPI_ISL_429984 | 2020-04-02 |
| L220F | USA | EPI_ISL_429974 | 2020-03-29 |
| L220F | USA | EPI_ISL_429977 | 2020-03-27 |
| L220F | USA | EPI_ISL_429980 | 2020-04-03 |
| L220F | USA | EPI_ISL_426465 | 2020-03-31 |
| L220F | USA | EPI_ISL_429975 | 2020-03-27 |
| L220F | USA | EPI_ISL_429979 | 2020-03-26 |
| L220F | USA | EPI_ISL_426454 | 2020-03-23 |
| L220F | USA | EPI_ISL_426458 | 2020-03-30 |
| L220F | USA | EPI_ISL_429983 | 2020-03-30 |
| L220F | USA | EPI_ISL_429986 | 2020-04-02 |
| L220F | USA | EPI_ISL_417517 | 2020-03-13 |
| L220F | USA | EPI_ISL_429976 | 2020-04-04 |
| L220F | USA | EPI_ISL_429978 | 2020-03-30 |
| L220F | USA | EPI_ISL_426456 | 2020-03-30 |
| L220F | USA | EPI_ISL_429985 | 2020-04-02 |
| L220F | USA | EPI_ISL_429973 | 2020-03-28 |
| L220F | USA | EPI_ISL_429972 | 2020-03-28 |
| L220F | USA | EPI_ISL_429969 | 2020-03-26 |
| **A191V/L220F** | **USA** |  |  |
| A191V/L220F | USA | EPI_ISL_426475 | 2020-04-04 |
| A191V/L220F | USA | EPI_ISL_419710 | 2020-03-12 |
| **N274D** | **France** | EPI_ISL_420610 | 2020-03-23 |
| **N274D** | **Finland** | EPI_ISL_418390 | 2020-03-13 |
| **M264I** | **USA** | EPI_ISL_427184 | 2020-03-25 |
| **A193V** | **USA** | EPI_ISL_427162 | 2020-03-30 |
| **N142S** | **USA** | EPI_ISL_427164 | 2020-03-31 |
| **C156F** | **USA** | EPI_ISL_430911 | 2020-04-08 |
| **L75F** | **USA** | EPI_ISL_421432 | 2020-03-17 |
| **L50F** | **Belgium** | EPI_ISL_434373 | 2020-03-28 |
| **R279C** | **France** | EPI_ISL_428365 | 2020-03-26 |
| **A191V** | **USA** | EPI_ISL_424172 | 2020-03-19 |
| **P132L** | **Russia** |  |  |
| P132L | Russia | EPI_ISL_428902 | 2020-03-23 |
| P132L | Russia | EPI_ISL_428892 | 2020-03-23 |
| **P132L** | **USA** | EPI_ISL_420579 | 2020-03-18 |
| **P168S** | **USA** | EPI_ISL_428789 | 2020-04-06 |
| **Q69H** | **USA** | EPI_ISL_426621 | 2020-04-01 |
| **P96L** | **USA** | EPI_ISL_427482 | 2020-04-01 |
| **V157L** | **USA** | EPI_ISL_426028 | 2020-03-04 |
| **L89F** | **Iceland** | EPI_ISL_424491 | 2020-03-20 |
| **L89F** | **USA** |  |  |
| L89F | USA | EPI_ISL_423028 | 2020-03-18 |
| L89F | USA | EPI_ISL_424260 | 2020-03-22 |
| L89F | USA | EPI_ISL_417376 | 2020-03-15 |
| L89F | USA | EPI_ISL_434526 | 2020-03-26 |
| L89F | USA | EPI_ISL_434525 | 2020-03-25 |
| L89F | USA | EPI_ISL_434530 | 2020-03-31 |
| L89F | USA | EPI_ISL_418036 | 2020-03-13 |
| L89F | USA | EPI_ISL_428335 | 2020-03-27 |
| L89F | USA | EPI_ISL_418893 | 2020-03-13 |
| L89F | USA | EPI_ISL_434517 | 2020-03-23 |
| L89F | USA | EPI_ISL_424868 | 2020-03-09 |
| L89F | USA | EPI_ISL_426627 | 2020-04-06 |
| **T45I** | **USA** |  |  |
| T45I | USA | EPI_ISL_421316 | 2020-03-23 |
| T45I | USA | EPI_ISL_421312 | 2020-03-20 |
| **A116V** | **USA** | EPI_ISL_427274 | 2020-03-20 |
| **A129V** | **USA** | EPI_ISL_430952 | 2020-03-28 |
| **P108S** | **Wales** |  |  |
| P108S | Wales | EPI_ISL_432197 | 2020-04-04 |
| P108S | Wales | EPI_ISL_432383 | 2020-04-05 |
| P108S | Wales | EPI_ISL_422042 | 2020-03-28 |
| P108S | Wales | EPI_ISL_432233 | 2020-04-01 |
| P108S | Wales | EPI_ISL_432300 | 2020-04-07 |
| P108S | Wales | EPI_ISL_432219 | 2020-04-04 |
| P108S | Wales | EPI_ISL_421005 | 2020-03-23 |
| P108S | Wales | EPI_ISL_432305 | 2020-04-07 |
| P108S | Wales | EPI_ISL_432275 | 2020-04-08 |
| P108S | Wales | EPI_ISL_422124 | 2020-03-29 |
| P108S | Wales | EPI_ISL_432309 | 2020-04-07 |
| **P108S** | **USA** | EPI_ISL_428787 | 2020-04-06 |
| **A266V** | **Australia** |  |  |
| A266V | Australia | EPI_ISL_419943 | 2020-03-21 |
| A266V | Australia | EPI_ISL_427683 | 2020-03-23 |
| A266V | Australia | EPI_ISL_427775 | 2020-03-25 |
| A266V | Australia | EPI_ISL_427682 | 2020-03-23 |
| A266V | Australia | EPI_ISL_427764 | 2020-03-23 |
| A266V | Australia | EPI_ISL_426725 | 2020-03-25 |
| A266V | Australia | EPI_ISL_426860 | 2020-03-27 |
| A266V | Australia | EPI_ISL_426862 | 2020-03-27 |
| A266V | Australia | EPI_ISL_426729 | 2020-03-26 |
| **A266V** | **USA** | EPI_ISL_421400 | 2020-03-17 |
| **A266V** | **USA** | EPI_ISL_421599 | 2020-03-21 |
| **A266V** | **USA** | EPI_ISL_421626 | 2020-03-21 |
| **A266V** | **Australia** |  |  |
| A266V | Australia | EPI_ISL_420012 | 2020-03-24 |
| A266V | Australia | EPI_ISL_419940 | 2020-03-21 |
| A266V | Australia | EPI_ISL_419938 | 2020-03-21 |
| **A266V** | **Scotland** | EPI_ISL_425911 | 2020-03-23 |
| **A266V** | **USA** | EPI_ISL_421348 | 2020-03-17 |
| **A266V** | **USA** | EPI_ISL_421615 | 2020-03-20 |
| **A266V** | **Jordan** | EPI_ISL_429992 | 2020-03-22 |
| **A266V** | **Australia** | EPI_ISL_426797 | 2020-03-24 |
| **A266V** | **Australia** | EPI_ISL_426771 | 2020-03-23 |
| **K236R** | **USA** |  |  |
| K236R | USA | EPI_ISL_434139 | 2020-03-27 |
| K236R | USA | EPI_ISL_434209 | 2020-03-31 |
| K236R | USA | EPI_ISL_434148 | 2020-03-27 |
| K236R | USA | EPI_ISL_434131 | 2020-03-27 |
| K236R | USA | EPI_ISL_434147 | 2020-03-27 |
| K236R | USA | EPI_ISL_434150 | 2020-03-27 |
| K236R | USA | EPI_ISL_434149 | 2020-03-28 |
| K236R | USA | EPI_ISL_434144 | 2020-03-27 |
| K236R | USA | EPI_ISL_434138 | 2020-03-27 |
| K236R | USA | EPI_ISL_434142 | 2020-03-27 |
| K236R | USA | EPI_ISL_434141 | 2020-03-29 |
| K236R | USA | EPI_ISL_426097 | 2020-03-25 |
| K236R | USA | EPI_ISL_427212 | 2020-03-27 |
| K236R | USA | EPI_ISL_424166 | 2020-03-19 |
| **A266V** | **Australia** | EPI_ISL_427808 | 2020-03-24 |
| **A266V** | **Australia** | EPI_ISL_427780 | 2020-03-23 |
| **A266V** | **Australia** | EPI_ISL_427679 | 2020-03-22 |
| **A266V** | **Australia** | EPI_ISL_427783 | 2020-03-25 |
| **A266V** | **USA** |  |  |
| A266V | USA | EPI_ISL_427472 | 2020-04-01 |
| A266V | USA | EPI_ISL_430414 | 2020-04-10 |
| A266V | USA | EPI_ISL_430332 | 2020-03-30 |
| A266V | USA | EPI_ISL_421733 | 2020-03-18 |
| A266V | USA | EPI_ISL_418198 | 2020-03-17 |
| A266V | USA | EPI_ISL_428762 | 2020-04-03 |
| A266V | USA | EPI_ISL_424939 | 2020-04-01 |
| A266V | USA | EPI_ISL_420572 | 2020-03-17 |
| **A266V** | **Australia** | EPI_ISL_419824 | 2020-03-21 |
| **A266V** | **Australia** | EPI_ISL_427794 | 2020-03-23 |
| **A266V** | **Australia** | EPI_ISL_427760 | 2020-03-24 |
| **A266V** | **Australia** | EPI_ISL_427787 | 2020-03-26 |
| **A266V** | **Australia** | EPI_ISL_427669 | 2020-03-22 |
| **A266V** | **Australia** | EPI_ISL_427688 | 2020-03-22 |
| **A266V** | **USA** |  |  |
| A266V | USA | EPI_ISL_427612 | 2020-03-16 |
| A266V | USA | EPI_ISL_427601 | 2020-03-16 |
| **A266V** | **Australia** |  |  |
| A266V | Australia | EPI_ISL_421636 | 2020-03-26 |
| A266V | Australia | EPI_ISL_427742 | 2020-03-21 |
| **A266V** | **Australia** | EPI_ISL_427785 | 2020-03-23 |
| **A266V** | **Australia** | EPI_ISL_427778 | 2020-03-25 |
| **A266V** | **Australia** | EPI_ISL_427782 | 2020-03-25 |
| **A266V** | **Australia** | EPI_ISL_427768 | 2020-03-23 |
| **A266V** | **Australia** |  |  |
| A266V | Australia | EPI_ISL_419956 | 2020-03-21 |
| A266V | Australia | EPI_ISL_427095 | 2020-04-03 |
| A266V | Australia | EPI_ISL_427094 | 2020-04-03 |
| A266V | Australia | EPI_ISL_427092 | 2020-04-03 |
| A266V | Australia | EPI_ISL_427689 | 2020-03-22 |
| **A266V** | **Australia** | EPI_ISL_427791 | 2020-03-24 |
| **A266V** | **Australia** | EPI_ISL_427708 | 2020-03-21 |
| **A266V** | **Australia** | EPI_ISL_419942 | 2020-03-21 |
| **A266V** | **USA** |  |  |
| A266V | USA | EPI_ISL_426052 | 2020-03-30 |
| A266V | USA | EPI_ISL_426056 | 2020-03-30 |
| A266V | USA | EPI_ISL_430430 | 2020-04-13 |
| A266V | USA | EPI_ISL_430927 | 2020-04-04 |
| A266V | USA | EPI_ISL_430921 | 2020-04-06 |
| A266V | USA | EPI_ISL_424271 | 2020-03-22 |

